# On the physiological and structural contributors to the overall balance of excitation and inhibition in local cortical networks

**DOI:** 10.1101/2023.01.10.523489

**Authors:** Farshad Shirani, Hannah Choi

## Abstract

Overall balance of excitation and inhibition in cortical networks is central to their functionality and normal operation. Such orchestrated co-evolution of excitation and inhibition is established through convoluted local interactions between neurons, which are organized by specific network connectivity structures and are dynamically controlled by modulating synaptic activities. Therefore, identifying how such structural and physiological factors contribute to establishment of overall balance of excitation and inhibition is crucial in understanding the homeostatic plasticity mechanisms that regulate the balance. We use biologically plausible mathematical models to extensively study the effects of multiple key factors on overall balance of a network. We characterize a network’s baseline balanced state by certain functional properties, and demonstrate how variations in physiological and structural parameters of the network deviate this balance and, in particular, result in transitions in spontaneous activity of the network to high-amplitude slow oscillatory regimes. We show that deviations from the reference balanced state can be continuously quantified by measuring the ratio of mean excitatory to mean inhibitory synaptic conductances in the network. Our results suggest that the commonly observed ratio of the number of inhibitory to the number of excitatory neurons in local cortical networks is almost optimal for their stability and excitability. Moreover, the values of inhibitory synaptic decay time constants and density of inhibitory-to-inhibitory network connectivity are critical to overall balance and stability of cortical networks. However, network stability in our results is sufficiently robust against modulations of synaptic quantal conductances, as required by their role in learning and memory.

**Summary:** We leverage computational tractability of a biologically plausible conductance-based meanfield model to perform a comprehensive bifurcation and sensitivity analysis that demonstrates how variations in key synaptic and structural parameters of a local cortical network affect network’s stability and overall excitation-inhibition balance. Our results reveal optimality and criticality of baseline biological values for several of these parameters, and provide predictions on their effects on network’s dynamics which can inform identifying pathological conditions and guide future experiments.

## Introduction

Stability and excitability are essential properties of cortical networks that are established through complicated dynamic interactions between neurons. A cortical network of one cubic millimeter in volume in mammalian neocortex is composed of tens of thousands of neurons. Each of these neurons receive excitatory and inhibitory synaptic inputs from over a thousand other neurons, both through long-range corticocortical fibers coming from neurons residing outside the network and through intracortical fibers coming from the neurons inside the network [1], [2, Figure 1]. Within the network, neurons are highly interconnected via all types of excitatory-to-excitatory, excitatory-to-inhibitory, inhibitory-to-excitatory and inhibitory-to-inhibitory connections. Such a massively interconnected network of dynamically interacting neurons must have intrinsic mechanisms to control the level of overall excitation and inhibition that is generated in the network at every instance of time. If the recurrent excitation provided by the population of excitatory neurons on itself—which is necessary for self-sustaining activity of the network—is not sufficiently balanced by the inhibition it receives from inhibitory neurons, then the overall level of excitation in the network can rapidly rise to an extreme level at which the spiking rates of the neurons saturate. Oppositely, if the inhibitory neurons impose excessive inhibition on the excitatory population, the network loses the level of excitability that it needs to effectively respond to inputs coming from other cortical areas. Hence, maintaining an overall balance of excitation and inhibition is crucial for the functionality of a cortical network.

**Figure 1:**
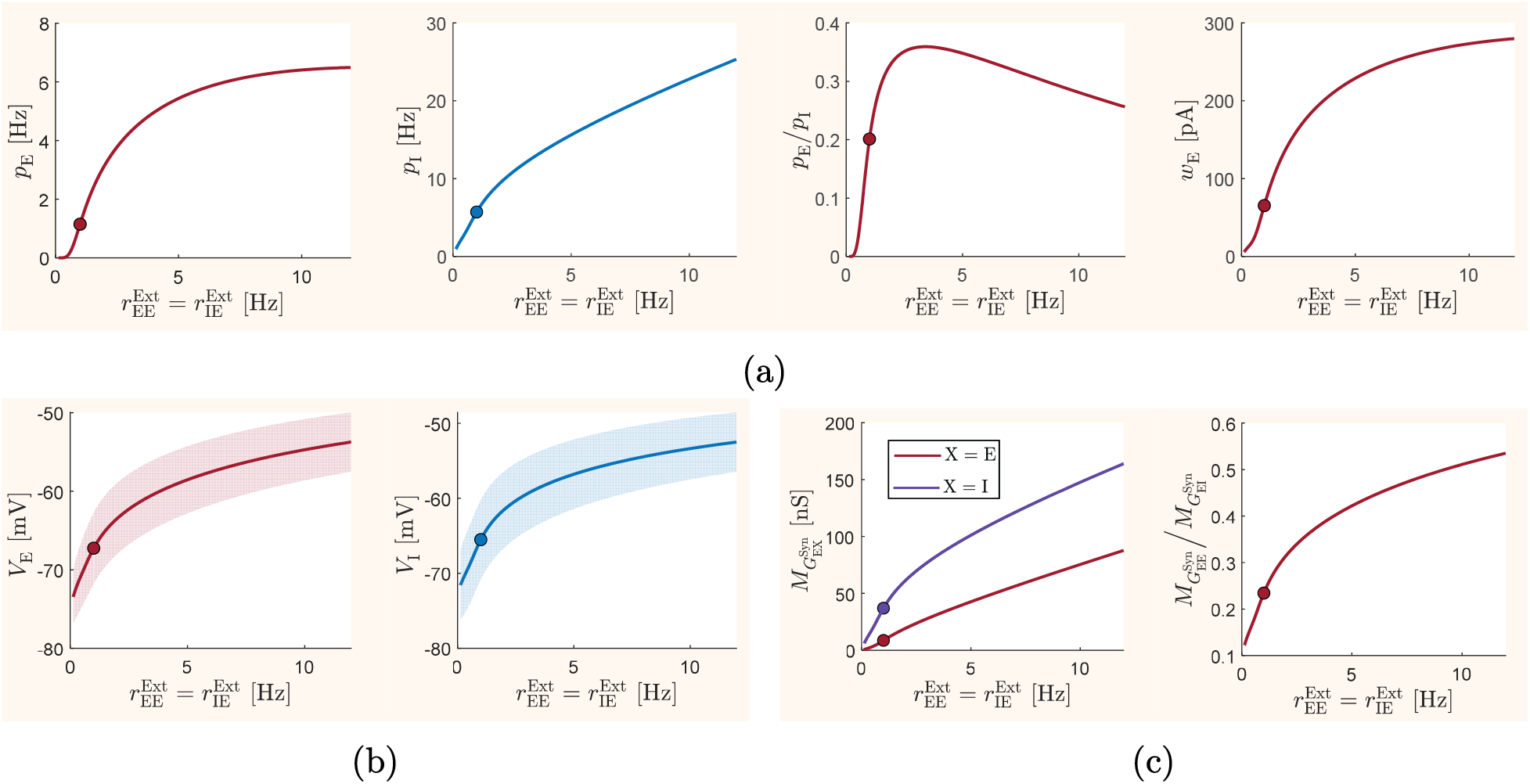
Steady-state mean-field activity with respect to variations in the mean frequency of the external inputs. All parameter values of the mean-field model are set to their baseline values given in Table 1. The model is driven by external inputs of different mean frequency 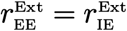, and the resulting steady-state values of different network quantities are shown in the graphs. The points marked by dots in the graphs correspond to the baseline mean input frequency of 1 Hz, which we considered as the level of background input to the mean-field model. (a) Mean excitatory firing rate *p*_e_, mean inhibitory firing rate *p*_i_, ratio between the mean firing rates *p*_e_/*p*_i_, and the mean excitatory adaptation current *w*_e_. (b) Excitatory membrane potential *V*_e_ and inhibitory membrane potential *V*_i_ of the neurons. Solid lines indicate the mean values *M*_*V*_e__ and *M*_*V*_i_ of the membrane potentials, and shaded areas indicate variations in the membrane potentials within a range of one-standard deviation (*S*_*V*_e___ and *S*_*V*_i__) from the mean values. (c) Mean synaptic conductances 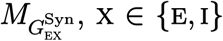 of the excitatory population, and the ratio between the two conductances. Mean synaptic conductances of the inhibitory population take the same values as those of the neurons of the excitatory population.

Theoretical and experimental studies have confirmed the existence of a dynamically regulated balance of excitation and inhibition in local cortical networks at multiple states of wakefulness and sleep [3–18]. It has been hypothesized that the balance of excitation and inhibition is essential for controlling network-level information transmission [19,20], efficient, high-precision, and high-dimensional representations and processing of sensory information [5, 11,13], enabling cortical computations by enhancing the range of network sensitivity to sensory inputs [8], selective amplification of specific activity patterns in unstructured inputs [21], maintaining information in working memory [22], and, importantly, preserving network stability [10, 23]. Pathological conditions resulting in deviations from normal levels of excitation-inhibition balance, hence hypo-or hyper-excitation in cortical networks, have been associated with several neurological disorders, such as Autism Spectrum Disorders, schizophrenia, mood disorders, Alzheimer’s disease, Rett Syndrome, and epilepsy [7, 9, 19, 23–27]. Nevertheless, neuromodulatory mediated deviations from finely balanced network states, as long as they do not result in pathological dysfunction, are also thought to be important in enabling certain network operations, such as performing longterm changes in sensory receptive fields [8,9,24].

There is, however, a broad range of interpretations of excitation-inhibition balance in the literature. For instance, *loose* balance of excitation and inhibition in a network has been defined as a state during which temporal variations in excitatory and inhibitory input currents to neurons are correlated on a slow time scale, whereas on a faster time scale they exhibit uncorrelated fluctuations. When faster fluctuations are also strongly correlated, with possibly a small time lag between them, the balance has been called *tight* [11,28,29]. From a related but mostly spatial point of view, *global* balance has been referred to a state at which each neuron of the network receives approximately equal amounts of excitation and inhibition, so that on average the overall levels of excitation and inhibition in the network are the same. The balance is called *fine-scale* or *detailed* when, in addition to having global balance, the amounts of excitation and inhibition to neurons also balance each other at finer spatial resolution, that is on subsets of synaptic inputs corresponding to specific signaling pathways [8,20,28]. The presence of such variety of interpretations has been an additional source of difficulty in developing techniques for experimentally measuring the excitation-inhibition balance in cortical networks and understanding its underlying regulatory mechanisms [16].

Due to experimental complexities and interpretational ambiguities, it is still not well-understood how the excitation-inhibition balance is established, what cellular and network properties are homeostatically adjusted to maintain it, and how it can be accurately and meaningfully measured [16,30,31]. Despite the challenges, however, numerous homeostatic mechanisms have been proposed as possible regulatory processes involved in maintaining the balance in cortical networks [8,10,19,23,28,30,32–35]. Moreover, it is convincing that the balance is most likely established and regulated locally, that means through internal recurrent interactions within a local cortical network, as global interactions between different regions of the cortex are predominantly excitatory and cannot be effectively balanced at a global scale by short-range activity of inhibitory neurons [2, 7, 11, 18, 36].

The goal of this paper is to leverage the computational tractability of a biologically plausible mean-field model to perform an extensive study on how variations in some of the key physiological factors—that control the kinetics of synapses—and key structural factors—that determine the overall topology of a cortical network—affect the overall balance of excitation and inhibition in a local cortical network and potentially result in loss of stability and critical phase transitions in the dynamics of the network. Synaptic properties of a network can be dynamically adjusted through homeostatic plasticity mechanisms to compensate for changes in excitatory and inhibitory activity in the network, and thereby regulate the network balance. Structural organization of a network determines the types and amounts of interactions between neuronal populations, and hence is central to establishing the overall balance of activity in the network. Hence, knowing how changes in each of the key synaptic and structural parameters of a network affect its overall balance, and whether or not such changes can create a critical state in the dynamics of the network, is important in understanding the homeostatic mechanisms that regulate the balance, and in identifying the sources of pathological conditions that may arise during cortical development or as a result of neurological diseases. We specifically demonstrate the effects of changes in physiological parameters such as synaptic decay time constants, synaptic quantal conductances, and synaptic reversal potentials, as well as structural parameters such as the ratio of the number of inhibitory neurons to the number of excitatory neurons, overall connectivity density of the network, and the density of inhibitory-to-inhibitory connectivity. Performing such an extensive study experimentally is not practical, nor is it using biologically detailed neuronal network models, elucidating our choice of a biologically plausible mean-field model in this study. The computationally affordable framework of our study allows for testing effects of fine modulations of a range of key parameters, providing predictions that can hint future experiments.

Our interpretation of a balanced state of excitation and inhibition is in the sense of its functionality. We characterize a balanced state as a network-level operational state that satisfies certain qualitative properties that are often reported in normally functioning networks. Specifically, we consider a (well-) balanced state as a state at which (1) the spontaneous activity of the neurons in the network are asynchronous and irregular, (2) the network is sufficiently excitable, (3) the spontaneous and stimulated activity in the network remain stable, and (4) the network responds rapidly to a reasonably wide range of external stimuli. We use the term “overall” balance to refer to such network-level balance, a term also alternatively used in the literature for global balance. The asynchronous and irregular network activity under this balanced state is commonly observed in globally and tightly balanced networks [7,10,11,13,15,20,29,37]. The neurons in such balanced state are mostly expected to be depolarized near their spiking threshold [3, 38]. We use the ratio of mean excitatory to mean inhibitory synaptic conductances to measure the balance in a network, which is a measure used effectively in some experimental studies [3]. Our results show that this mean conductance ratio is a reliable measure to continuously quantify deviations from the balanced state, as it changes monotonically when the network balance is deviated from its reference value due to variations in the network parameters we study.

Our analysis is based on the biologically plausible mean-field model introduced in [39], with an additional neuronal adaptation mechanism proposed in [40]. This model, which has been developed in a sequence of works described in [39–43], has succeeded in fairly accurately predicting the mean spontaneous activity of neurons in a local cortical network during asynchronous irregular firing regimes, as well as their responses to certain external stimuli [39, 40]. We realistically characterize this model by setting the values of its biophysical parameters according to estimates obtained for cortical neurons of the mouse and rat brain [37, 44]. As a result, the model presents a balanced state of excitation and inhibition with mean firing activity of the neurons being very close to that observed in biophysically detailed models of rat neocortical microcircuitry [37]. We use the long-term spontaneous mean-field activity predicted by this model, when driven by a constant rate of background input spikes, to make our observations on the level of balance in the network. The mean-field framework of the model allows us to employ standard numerical bifurcation analysis techniques to investigate how the overall (mean-field) balance of the network, established at baseline parameter values, is affected by continuous changes in each of the physiological and structural network parameters over a wide range of biologically plausible values. In particular, we observe that in most cases the parameter changes that result in over-excitation in the network can eventually become critical and lead to a phase transition in the network activity to a high-amplitude slow oscillatory bursting regime. We partially verify some of the key predictions of our mean-field-based analysis using a more detailed spiking neuronal network model, as well as in comparison with some experimental and computational results available in the literature on the rat somatosensory microcircuitry [37].

We organize this paper as follows. In the Methods section, we provide an overview of our bifurcation and sensitivity analysis framework. In the Model Description section, we describe the details of the models we use to perform our analyses. In the Results section, we present the computational results we obtain based on the model. Finally, in the Discussion section, we summarize and discuss the key observations we make in our study. The results presented in the Results section are modular. All bifurcations diagrams we obtain for different biophysical quantities of the network are included in a single figure, provided separately for each of the physiological and structural parameters we study. Therefore, an alternative quick read can be made by skipping the entire Results section at first and proceeding directly to the Discussion section. The details for each of the key observations summarized in the Discussion section can then be found back in the corresponding Figures and their descriptions provided in the Results section.

## Methods

Our approach in studying the sensitivity of the overall balance of a network to continuous variations in network parameters relies on computing changes and phase transitions in longterm mean-field activity of the network. In the absence of sensory or cognitive stimuli, the activity of a local cortical network *in vitro* is driven mainly by *background* spikes from neighboring cortical areas. However, the mean rate of such background input spikes is low. Therefore, the spontaneous activity of an unstimulated local network can be expected to be predominantly self-generated. Assuming that the network is well-balanced, this spontaneous spiking network activity is asynchronous and irregular. Assuming further that the mean rate of background input spikes to the network is constant—which is a reasonable assumption we make in our analyses to be able to clearly distinguish the effects of parameter variations in our observations—then the mean spontaneous firing activity of the balanced network reaches quickly to a steady-state. Therefore, measuring the long-term (steady-state) mean-field activity of the network, driven by a constant rate of background spikes, provides good estimates for measuring the overall balance of excitation and inhibition in the network.

We use a conductance-based mean-field model, whose details are described below in the Model Description section, to compute approximate mean-field activity of a local cortical network. We set the parameter values of this model equal to the realistic values estimated for cortical networks in the mouse and rat brain. We begin our analyses by first verifying that the model with these preset parameter values, when driven by a realistic rate of background input spikes, presents mean-field activity consistent with the activity observed in a well-balanced state—as we described in our interpretation of overall balance in the Introduction section. We use the steady-state (equilibrium) mean-field activity computed at this state as a reference for balanced network activity. As our results show, the ratio of the mean excitatory to inhibitory synaptic conductances in the network is a reliable measure of overall balance in the network. Therefor, we use the value of this ratio, computed at the steady-state of the balanced network, as a reference for our quantification of deviations from the balanced state.

We employ numerical bifurcation analysis techniques to predict how the network balance is deviated from its reference state when we continuously vary the key physiological and structural parameters of the network. We individually study the effect of variations in each network parameters by considering it as a bifurcation parameter in a codimension-one continuation of the stable network equilibrium, that is the steady-state mean-field activity associated with the reference balanced state we described above. For each study, we demonstrate how the steady-state values of important network quantities, such as mean neuronal firing rates, mean membrane conductances, mean membrane potentials, and mean synaptic currents change as the bifurcation parameter varies within its entire range of biologically plausible values. Moreover, when a phase transition to an oscillatory behavior is detected at a bifurcation point on the curves of equilibria, we additionally perform codimension-one continuation of the emerging limit cycles. This allows us to observe how the frequency of the oscillations changes with respect to variations in the bifurcation parameters. We perform our bifurcation analyses using MatCont, version 7.3, [45], and individually report the results of each study in the Results section.

In our codimension-one analyses of the effects of synaptic parameters, we only consider variations in inhibitory synaptic parameters. However, we additionally perform codimension-two continuation of the network equilibrium by simultaneously considering both excitatory and inhibitory synaptic parameters as bifurcation parameters. This allows us to observe how joint modulation of the kinetics of excitatory and inhibitory synaptic activity affects the overall balance of excitation and inhibition in the network. We perform similar analyses to also investigate certain joint contribution of synaptic and structural factors.

Although utilizing the simplicity of a mean-field model enables us to perform an extensive study on the effects of variations in multiple network parameters on the overall network balance, the generality of our observations can suffer from the inevitable simplifying assumptions of the mean-field model construction. To address this concern, we also construct a spiking neuronal network model with equivalent structural, synaptic, and cellular parameters to those in the mean-field model. We then use this model to partially verify that some of the key predictions of our study based on the mean-field model still remain qualitatively valid when the simplifying assumptions of the mean-field description are removed. For this, we simulate the spiking neuronal network for two different sets of parameter values based on the predictions of the mean-field model, one corresponding to the network’s reference balanced state with asynchronous and irregular neuronal firing activity, and the other corresponding to an imbalanced oscillatory bursting state. We compare the results obtained from the two models using the same quantitative measures as those previously computed for the mean-field model.

Before presenting the results of our study, we provide below the detailed description of the mean-field and spiking neuronal network models, including the equations we use to compute the important descriptive quantities of the network.

### Model Description

Our computational study approach, as described above, relies on mathematical models whose detailed description, along with a discussion on the choice of their realistic parameter values, is provided below. The results we obtain based on these models are given in the Results section. Our key observations and their biological implications are discussed in the Discussion section.

#### Mean-field neuronal population model

The mean-field model we use here has been developed based on the Markovian master equations provided in [41], which describe the overall activity of a randomly connected balanced network of neurons in an asynchronous irregular spiking regime. Specifically, these equations present the temporal evolution of the mean and variance of the firing rates of neurons within the excitatory and inhibitory populations of the network, as well as the covariance of the firing rates between the two populations. The master equations given in [41] have been extended in [40] by including an additional equation that allows for spike frequency adaptation of the neurons.

Application of the master equations given in [41] requires developing neuronal transfer functions that characterize the stationary firing rate of neurons in response to their stationary presynaptic excitatory and inhibitory spiking activity. Such transfer functions provide population-level description of the neuronal activity and capture the specific properties of the single-neuron models and synaptic interactions that are considered in building the meanfield model of the entire network. Due to the nonlinearities involved in incorporating such properties, analytical derivation of the transfer functions is often quite challenging. Here, as in [40], we use the semi-analytically calculated transfer functions that are proposed in [42] and [43] under the assumption that the characterization of the transfer functions depends only on the statistical properties of subthreshold membrane potential fluctuations. These transfer functions rely on an effective membrane potential threshold for each excitatory and inhibitory population. This effective threshold is expressed as a second-degree polynomial on the moments of the subthreshold membrane potentials within each population, namely, on the mean, standard deviation, and autocorrelation time constant of the membrane potential fluctuations. The coefficients of this second-degree polynomial are obtained by fitting it to the dynamics of numerically simulated single neurons at different excitatory and inhibitory presynaptic spiking frequencies [40, 42, 43].

Below, we briefly provide the formulation of the mean-field model as given in [40], with several notational changes, some modifications for incorporating external inputs to the network, and a correction on the equations governing neuronal adaptation.

#### Mean-field model equations

To present the equations of the mean-field model, let e and i denote, respectively, the excitatory and inhibitory neuronal populations of a local cortical network composed of a total number of N neurons. Note that N = N_e_ + N_i_, where N_e_ and N_i_ denote the total number of excitatory and inhibitory neurons in the network, respectively. For all time *t* ∈ [0,*T*], *T* > 0, and population types x and y, where x,y ∈ {e, i}, the modeled neuronal activity is represented by the following variables:

- *p*_x_(*t*), measured in Hz, denoting the mean firing rate of neurons in the x population at time *t*,
- *q*_xy_(*t*), measured in Hz^2^, denoting the covariance of the firing rates between the x and y populations at time *t*,
- *w*_x_(*t*), measured in pA, denoting the mean adaptation current of neurons in the x population at time *t*,
- 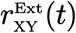, measured in Hz, denoting the average rate of spikes received by neurons of x population at time *t* through each of the afferent fibers arriving from external neurons of type y. Although in general some fraction of the afferent fibers can arrive from external inhibitory neurons, throughout this paper we assume all these external input spikes to the network are received only from excitatory neurons.

The system of differential equations that governs the time evolution of the state variables *p*_x_, *q*_xy_, and *w*_x_ for a given external drive 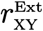 are provided by the master equations (13)–(15) below. Note that *q*_ei_ = *q*_ie_, hence it is sufficient to solve (13)–(15) only for one of these two quantities. As stated above, these master equations require calculation of the transfer functions that relate the firing rate of neurons in each population to their presynaptic excitatory and inhibitory spike rates. The conductance-based internal interactions that yield the derivation of these transfer functions are modeled as follows [39, 42, 43].

Let 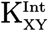, with x, y ∈ {e, i}, denote the average number of presynaptic connections that neurons within the x population in the network receive internally from the neurons of the y population of the network. Similarly, let 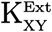 denote the average number of presynaptic connections that neurons within the x population receive from external neurons of type y residing outside the network. Then, the average number of presynaptic connections that neurons within the x population receive in total from both internal and external neurons of type y is given as

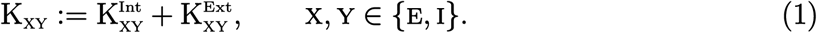

As stated above, throughout this paper we assume all external cortical connections to the local network under the study are of excitatory type, that is, 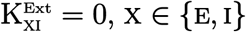. Moreover, letting P_xy_ denote the connection probabilities between neurons of x and y populations, as described in Table 1, the average number of internal connections are given as

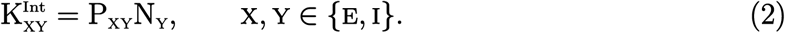

**Table 1:**
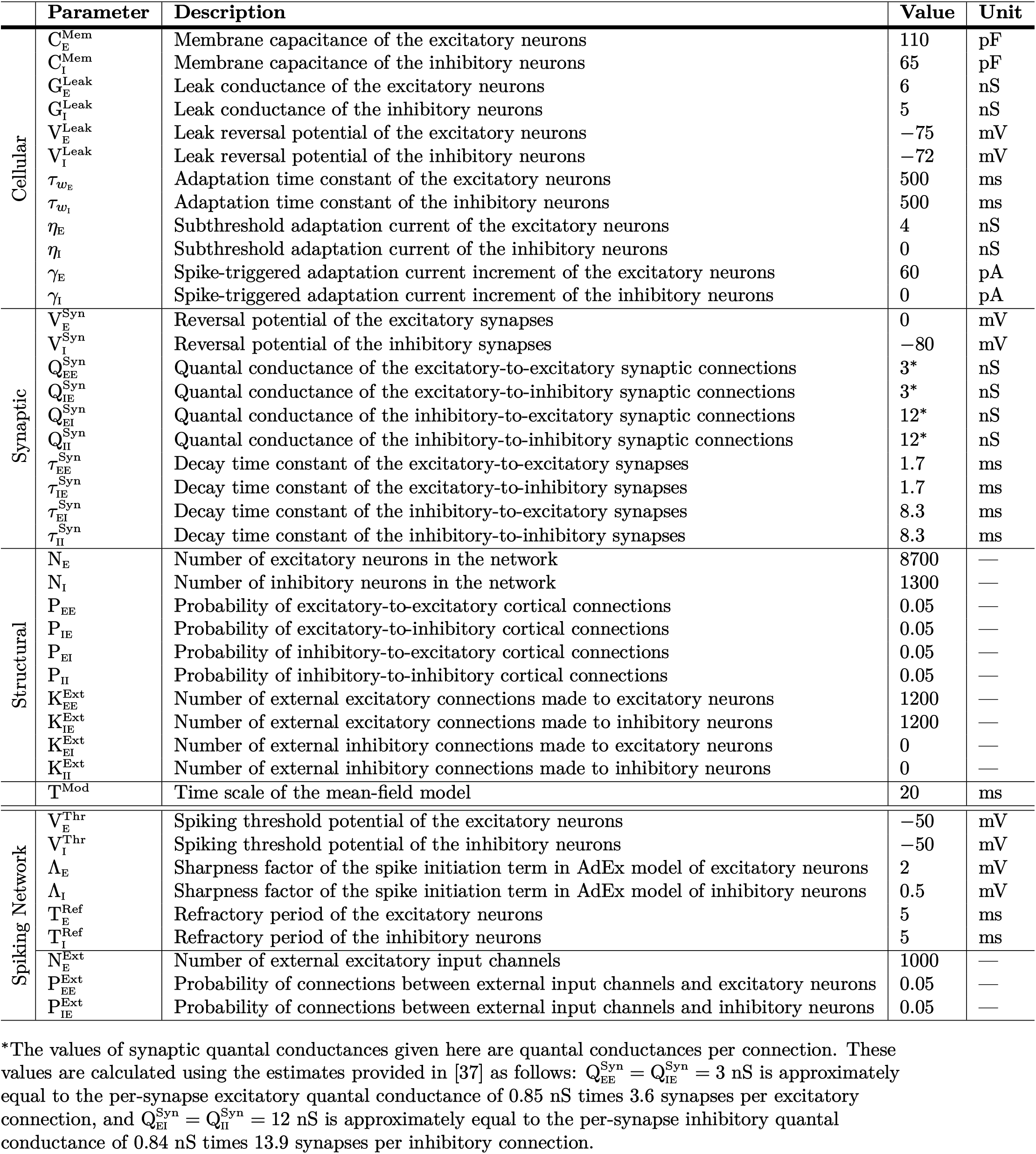
Cellular, synaptic, and structural parameters of the mean-field and spiking network models.

Next, let *r*_xe_ and *r*_xi_ denote, respectively, the average rate of excitatory and inhibitory presynaptic spikes that neurons in a population of type x receive both internally from other neurons within the network and externally from neurons residing outside the network. That is,

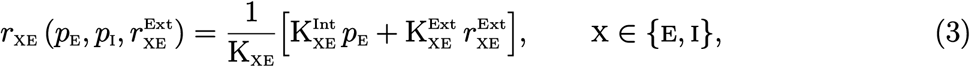

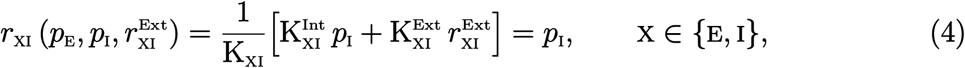

where the second equality for *r*_xi_ is due to the assumption 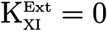.

The train of spikes received by a neuron from its presynaptic neurons dynamically change the neuron’s membrane conductance. Let 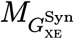 and 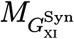 denote, respectively, the mean value of the total excitatory and inhibitory synaptic conductances of a neuron of type x. Note that, throughout the paper, we will use *M_A_* and *S_A_* to denote the mappings that give the mean and standard deviation of the quantity *A*, respectively. The total excitatory/inhibitory synaptic conductance of a neuron of type x is the conductance resulting cumulatively from all the synapses that are made on this neuron by other excitatory/inhibitory neurons in the network. The effective value of these synaptic components depend on the rate of input spikes to the neurons. As in [46] and [43], the mean conductances can approximately be given as

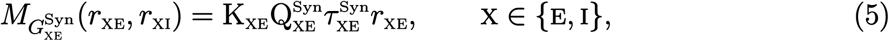

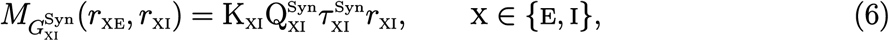

which then allow for the calculation of *M*_*G*_x__, the mean value of the total membrane conductance of the neurons within the x population, as

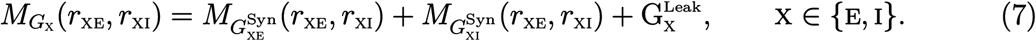

The parameter 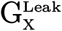 in (7) denotes the leak conductance of the neurons. The parameters 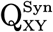 and 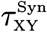 in (5) and (6), with x, y ∈ {e, i}, denote the quantal conductance and decay time constant of the synaptic connections, as described in Table 1. Note that these synaptic parameters correspond to the first-order model of synaptic conductance kinetics given by equation (18) below. Note also that neurons typically receive multiple synapses per single connection. The parameters 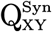 in (5) and (6) denote synaptic quantal conductances *per connection*, and hence they are equal to the sum of the per-synapse quantal conductances generated by each of the synapses of a neuronal connection.

An approximation of the mean membrane potential of the x population based on the mean conductances calculated above has been provided in [46] and [43] as

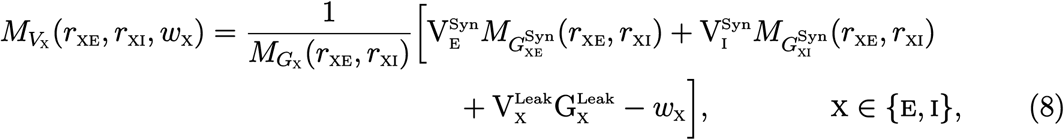

which can be roughly understood as a steady-state current law on the different currents flowing through the membrane of a neuron, with the neuron’s membrane potential and conductances being replaced with their mean values across the population. The parameters 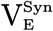 and 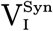 denote the excitatory and inhibitory synaptic reversal potentials as described in Table 1. We assume these parameters do not depend on the type of the postsynaptic neurons.

The following approximations for the standard deviation *S*_*V*_x__ and a global autocorrelation time constant (approximate speed) *T*_*V*_x__ of membrane potential fluctuations have been obtained in [42] and [43] as

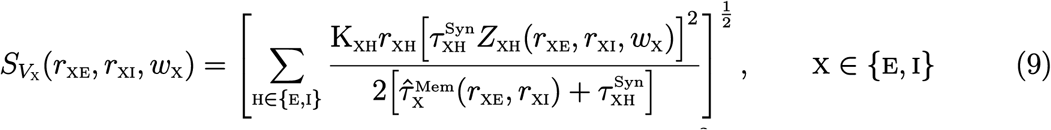

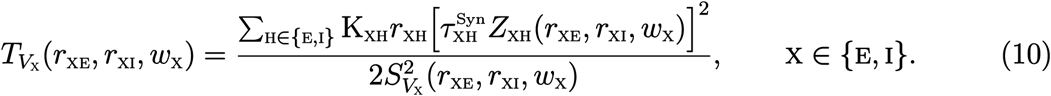

where

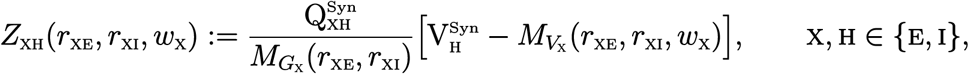

and 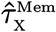 denotes the effective membrane time constant of the x population, given by

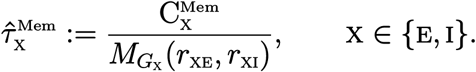

The parameter 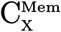 denotes the membrane capacitance of the neurons as described in Table 1.

The transfer functions *F*_x_ of neurons within each population x can now be characterized using the membrane potential moments *M*_*V*_x__, *S*_*V*_x__, and *T*_*V*_x__ given by (8), (9), and (10), respectively. For this, a semi-analytic form has been proposed in [42] as

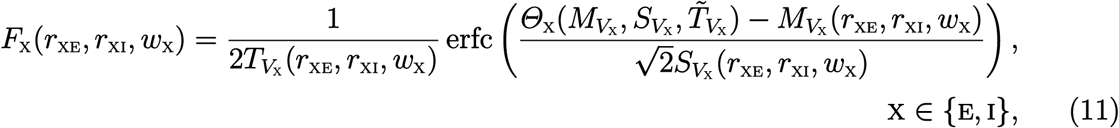

where erfc is the complementary error function and 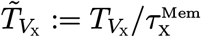, with 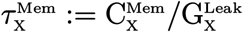 denoting the membrane time constant of the neurons of type x.

The effective membrane potential threshold *Θ*_x_ in (11), which accounts for the nonlinearities in the neuronal dynamics, is expressed in [40] as the following second-degree polynomial

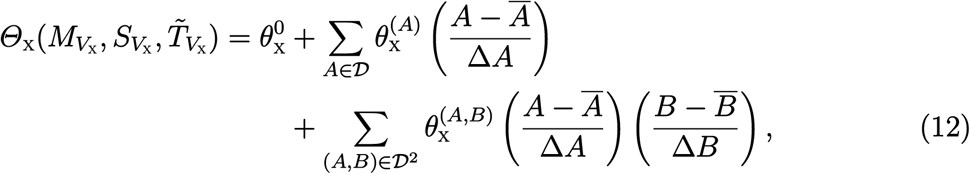

where 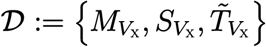 and 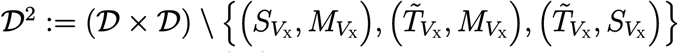. The normalization parameters 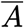 and 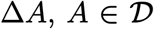, in (12) are not population type specific and their values are given in Table 2. It should be noted that some of these parameter values appear to be misreported in [40]. Here, we used their values as originally given in [42]. The coefficients 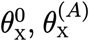, and 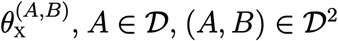, are given in Table 3. They are calculated in [40] by fitting the effective thresholds and the resulting transfers function to numerically calculated dynamics of adaptive exponential integrate and fire (AdEx) neurons. Note that the fit parameters in Table 3 are provided for both regular-spiking and fast-spiking neurons. Unless otherwise stated, we assume throughout this paper that all excitatory neurons are regular-spiking and all inhibitory neurons are fast-spiking.

**Table 2:** Normalization parameters of the semi-analytic transfer functions [42].

| Cell type | $\overline{M_{V_x}}$ | $\overline{S_{V_x}}$ | $\overline{\tilde{T}_{V_x}}$ | $\Delta M_{V_x}$ | $\Delta S_{V_x}$ | $\Delta \tilde{T}_{V_x}$ |
| --- | --- | --- | --- | --- | --- | --- |
| x = E, I | −60 mV | 4 mV | 0.5 | 10 mV | 6 mV | 1 |

Finally, calculation of the semi-analytic transfer functions (11) allows us to complete the presentation of the mean-field model by providing the masters equations that govern the evolution of the population-level neuronal activity in the network. For x ∈ {e, i}, y ∈ {e, i}, and *t* ∈ [0,*T*], *T* > 0, the equations are given as

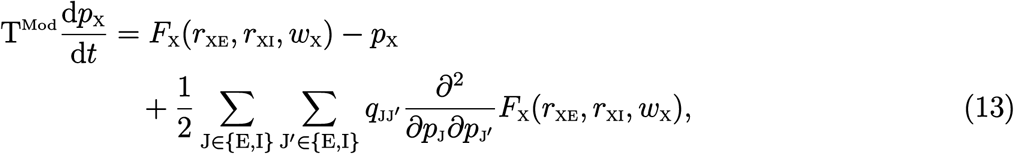

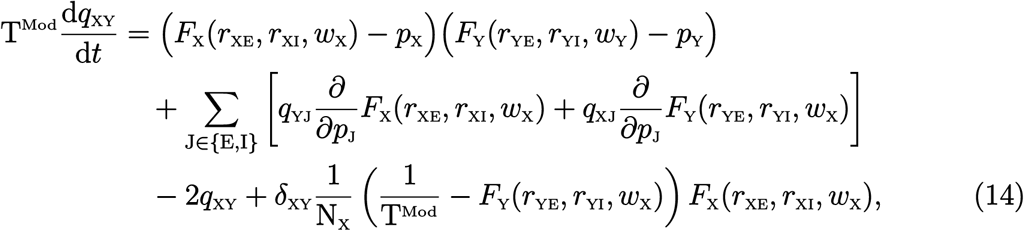

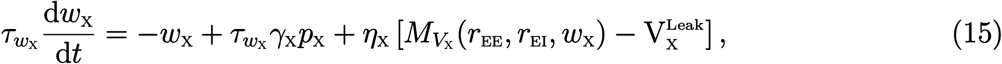

where T^Mod^ is the modeling time scale of the Markovian description of the network activity considered in [41], and *τ*_*w*_x__, *γ*_x_, and *η*_x_ are adaptation parameters of the neurons as described in Table 1. Moreover, *δ*_xy_ denotes the Kronecker delta function, that is, *δ*_xy_ = 1 if x = y, and *δ*_xy_ = 0 if x ≠ y. Note that, for simplicity of the exposition, the dependence of variables *p*_x_, *q*_xy_, and *w*_x_ on *t* is not explicitly shown in (13)–(15). Moreover, note that *r*_xy_ depends on the variables *p*_e_, *p*_i_ and the external inputs 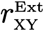, as in (3) and (4).

Equations (13) and (14), which give the temporal evolution of the mean and covariance of the firing rates have been developed in [41]. The additional equation (15), which captures the dynamics of sub-threshold and spike-triggered neuronal adaptation corresponding to the adaptation equation of a single-neuron AdEx model [47], has been provided in the set of equations in [40]. Note that (15) includes a correction on the original equation [40, Equ. (2.6)], as this equation suffers from a unit mismatch between the physical quantities and does not correctly correspond to the adaptation equation of the network’s constitutive AdEx neurons.

We numerically solve (13)–(15) and analyze bifurcations in the equilibrium solutions of these equations to obtain the solution curves and bifurcation diagrams presented in the Results section. The distinction between the dynamics of the excitatory and inhibitory neuronal populations is made through inclusion or exclusion of the adaptation equation (15) and corresponding choice of the fit parameters given in Table 3. We consider all excitatory neurons to be regular-spiking, meaning that their dynamics undergoes adaptation. Unless otherwise stated, we consider all inhibitory neurons to be fast-spiking, with no adaptation. Therefore, for the fast-spiking inhibitory population we set *w*_i_ = 0 and exclude (15) for x = i from the numerical computations.

**Table 3:**
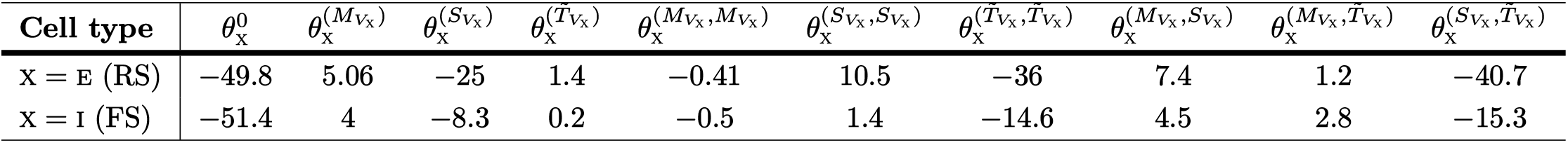
Fit parameters of the semi-analytic transfer functions for populations of regular-spiking (RS) and fast-spiking (FS) neurons [40].

In the numerical analysis presented in the Results section, we also investigate dynamic variations in the mean value of the synaptic currents flowing through the membrane of the neurons in each population. We use the approximate steady-state current law (8) to derive the following approximations:

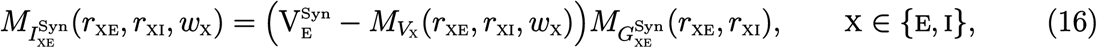

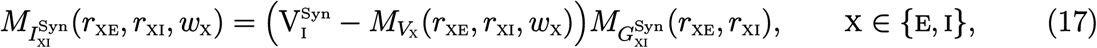

where 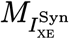 and 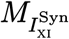 denote the mean excitatory and inhibitory synaptic currents in the x population, respectively.

#### Mean-field model parameters

The description and value of the biophysical parameters of the mean-field model presented above are given in Table 1. The parameter values are chosen to be in the range of realistic parameter values reported in the literature for the mouse and rat cortex. With these parameter values, which hereafter we refer to as *baseline parameter values*, the dynamics of the model shows a proper balance of excitation and inhibition wherein the mean firing rates of the excitatory and inhibitory populations closely coincide with those of the rat neocortex in an asynchronous irregular regime—as observed through both *in vivo* measurements and biophysically detailed *in silico* reconstruction of the rat neocortical microcircuitry [37]. Correspondingly, we refer to this state of balanced activity in the network as *baseline balanced state*.

The values of the passive membrane parameters 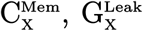, and 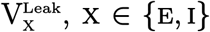, are chosen to be approximately equal to the median values given in [44, Suplementary Table 2], with the membrane leak conductance being reciprocal to the membrane resistance given in there. The range of values provided in [44] are obtained by tuning the parameters of a family of generalized leaky integrate-and-fire models so that they reproduce the spiking activity of a large number of recorded neurons in the primary visual cortex of adult mouse. The associated neuronal recordings are available in the Allen Cell Types Database [48].

The values of the adaptation parameters *τ*_*w*_e__, *η*_e_, and *γ*_e_ given in Table 1 are equal to the values chosen in [40] for these parameters. These values are used in [40] in the calculations resulting in the fit parameters given in Table 3 for regular-spiking excitatory neurons. Note that, as in [40], we assume all inhibitory neurons in the baseline model are non-adapting. Moreover, note that the values of the adaption parameters chosen here are also comparable to the values provided in [47] by fitting an AdEx model to dynamics of a biophysically detailed conductance-based model of a regular-spiking neuron.

The values of the synaptic reversal potentials given in Table 1 are fairly typical. We choose the baseline values for the other synaptic parameters to be equal to their mean values as given in [37]. The parameter values provided in [37] are used for a detailed digital reconstruction of the juvenile rat somatosensory microcircuitry as part of the Blue Brain Project [49]. Specifically, we set the values of 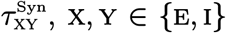, equal to the values given in [37, Table S6]. For synaptic quantal conductances, however, an adjustment is made on the values provided in [37]. The average quantal conductance value of 0.85 nS stated in [37, Page 471] for excitatory synapses, and the average value of 0.84 nS stated for inhibitory synapses, are specified as per-synapse conductances. Whereas, as stated in the model description above, the parameters 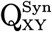 in the mean-field model we use here are associated with synaptic quantal conductances per connection. Therefore, to make adjustment for this difference, we scale the average quantal conductances in [37] by the average number of synapses per connection. These scaling values are also provided in [37, Page 464], as 3.6 synapses per connection for excitatory connections and 13.9 synapses per connection for inhibitory neurons. The adjusted conductances are then given in Table 1 as values of 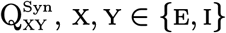.

We consider a randomly connected network composed of a total number of N = 10000 neurons, out of which N_e_ = 8700 neurons are excitatory and the remaining N_i_ = 1300 are inhibitory. This relatively large size of the network allows for the mean field approximation described above, and is also comparable to the size of the neuronal populations in layers 2/3, 5, and 6 of the rat neocortical microcircuitry investigated in [37]. The 87% excitatory versus 13% inhibitory proportions of the neurons are chosen according to the overall estimates provided in [37, Page 461].

Choosing the parameter values for internal and external connectivity requires some con-siderations and simulation-based adjustments. The local connectivity density of neuronal networks varies across cortical regions and layers. Moreover, more than 80% of synapses in a local network of nearly 200 micrometer in diameter come from external neurons residing outside of the network [37,50]. As a result, it is estimated that when a slice of cortical tissue with a typical thickness of 300 micrometer is cut from the cortex, only about 10% of excitatory synapses and about 38% of inhibitory synapses remain intact [50]. Therefore, the balance of excitation and inhibition in such cortical slice is largely deviated toward over-inhibition. Taking these into account, we choose the connectivity parameters of the mean-field model so that the resulting network, driven by a biologically reasonable rate of background spikes from external neurons, presents a balance of excitation and inhibition with mean excitatory and inhibitory firing rates being consistent with those measured in a detailed digital reconstruction of the rat cortical microcircuitry in an asynchronous irregular spiking regime [37].

We set the connection probability of all types of internal network connections to be equal to 0.05. For excitatory-to-excitatory connections, this value is quite comparable to the values reported in the literature for different cortical layers [37, 51, 52]. For other types of connections, this value appears to be almost half the typical values estimated experimentally. However, experimental estimations of neuronal connectivity are often obtained using cortical slices. Hence—due to the uneven reduction in the number of excitatory and inhibitory connections during slicing, as described above—using such estimates of inhibitory connection probabilities for a structurally simplified network such as the one we consider here can potentially result in an over-inhibited network. Our preliminary simulations of the meanfield model with larger values of inhibitory connection probabilities shows an imbalance of activity as expected, with the model requiring excessive amount of external excitatory drive in order to produce a reasonable firing activity. Therefore, to achieve a reasonable balance of activity, we choose inhibitory connection probabilities smaller than the experimentally obtained estimates.

We assume neurons of the network do not receive any external inhibitory inputs, whereas they each receive an average number of 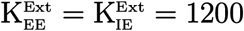 excitatory connections from external neurons. This number of external inputs is comparable with the estimates provided in [52, Table 3]. Moreover, with this number of external connections, along with the population size and internal connection probability chosen above, each neuron receives more than 60% of its synapses from external neurons, a percentage comparable to the estimates given in [37] and [50] as discussed above. Additionally, with the chosen number of external connections and internal connectivity, and with an average external (background) spiking rate of 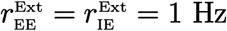, the numerical simulation of the model results in excitatory and inhibitory firing rates which are very close to those obtained in reconstructed rat microcircuitry [37]. The details of the simulations are provided in the Results section.

Finally, note that in order for the Markovian assumption used in the derivation of the mean-field model to be valid, the time scale of the model, T^Mod^, must be small enough so that neurons fire a maximum of one spike at each Markovian time step of length T^Mod^, and must be large enough so that the network dynamics can be considered memoryless over the timescale of T^Mod^; see [39,40,43]. Here, we set the value of T^Mod^ to be approximately equal to the time constant of the neurons, as suggested in [40] and [39]. This completes the discussion of the parameter values for the mean-field model. The rest of the parameters given at the bottom of Table 1 belong to the spiking network model described below.

### Spiking neuronal network model

We construct a network of randomly connected AdEx spiking neurons, in direct correspondence to the mean-field description presented above. For this, first let 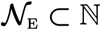 and 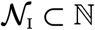, such that 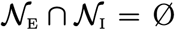, be two ordered sets that index neurons of the excitatory and inhibitory populations, respectively. Note that 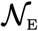 is of cardinality N_e_, and 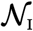 is of cardinality N_i_. The total number of neurons in the network is then indexed by the set 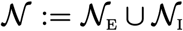. Also, let 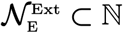, such that 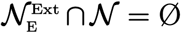, be an ordered set that indexes the external excitatory neurons that project onto at least one neuron inside the network. Note that, similar to the mean-field description, we assume no external inhibitory inputs to the network. With 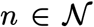, the neuronal activity of the spiking network is then represented by the following variables:

- *v_n_*(*t*), measured in mV, denoting the membrane potential of the *n*th neuron at time *t*,
- *w_n_*(*t*), measured in pA, denoting the adaptation current of the *n*th neuron at time *t*.

A neuron in the network fires a spike at a time *t* = *t*\* when its membrane potential exceeds its spiking threshold potential denoted by 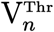, that is 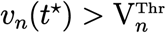. Let 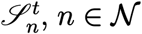, denote a set that stores all spike times of the *n*th neuron up to time *t*. Let, moreover, 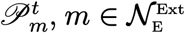, denote a set that stores all spike times of the *m*th excitatory external neuron up to time *t*. As described below, we assume these external spike times are Poisson-distributed.

Now, let *g_nm_*(*t*) denote the membrane synaptic conductance of the *n*th postsynaptic neuron generated through its synaptic connection with the *m*th presynaptic neuron. Let the set 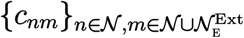 capture the connectivity of the network, including connections from external neurons, such that *c_nm_* = 1 if there is a connection from the *m*th neuron to the *n*th neuron, and *c_nm_* = 0 otherwise. Then, approximating kinetics of the synaptic conductance at each presynaptic spike time *t*\* by an instantaneous rise to a peak (quantal) conductance 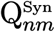 followed by an exponential decay with time constant 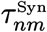, the membrane synaptic conductances are given as

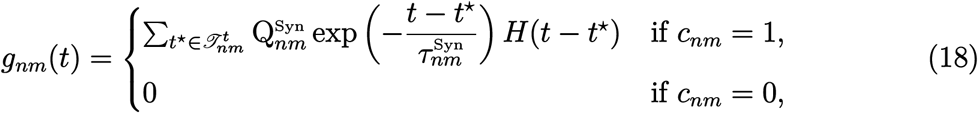

where *H* denotes the Heaviside step function and

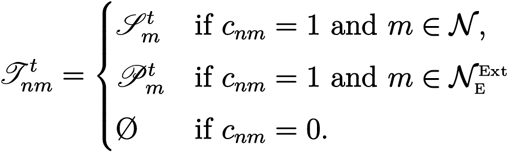

Next, the total synaptic current 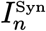 to the *n*th postsynaptic neuron can be computed as a function of *v_n_* and *t*, by adding together all fractions of current coming from each individual synapse that the neuron receives. Let 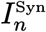 be decomposed into its excitatory and inhibitory components as 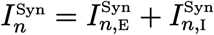. Then,

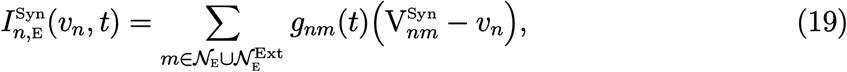

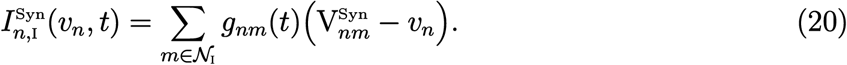

where 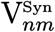 denotes the reversal potential of the synapse between the *n*th postsynaptic and *m*th presynaptic neurons when *c_nm_* = 1.

Finally, representing each neuron by an AdEx model [47], the subthreshold activity of the network is given by the following system of differential equations for all 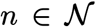 and *t* ∈ [0,*T*], *T*> 0,

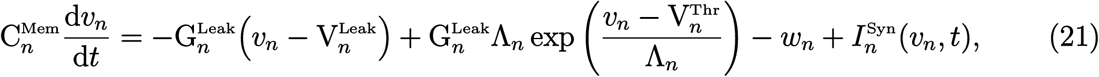

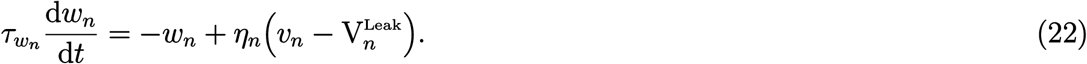

The parameters 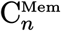 and 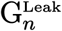, and 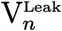 in (21) denote the membrane capacitance, leak conductance, and leak reversal potential of the *n*th neuron, respectively. Moreover, 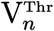 and Λ_*n*_ denote, respectively, the spiking threshold potential of the *n*th neuron and its sharpness factor for spike initiation. The parameters *τ_w_n__* and *η_n_* in (22) denote the adaptation time constant and the subthreshold adaptation current of the *n*th neuron, respectively.

The spiking activity of the network is captured by performing the following updates at each time instance *t* = *t*\* for all neurons that fire a spike at *t* = *t*\*, that is, for all 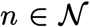 such that 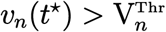:

- *v_n_* is reset to the resting potential, denoted by 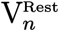, and is kept at this value for a duration of time equal to the refractory period of the *n*th neuron, 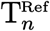,
- *w_n_* is incremented by a constant amount *γ_n_*,
- *t*\* is added to the set 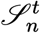.

For the numerical analysis presented in the Results section, we set the parameters of all excitatory neurons and synapses, as well as all inhibitory neurons and synapses, to be the same, and equal to the baseline parameter values given in Table 1. That is, for all 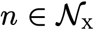 and x ∈ {e, i}, we set 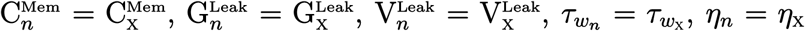, and *γ_n_* = *γ*_x_, in accordance with the baseline parameters used for the mean-field model. We also set the specific parameters of AdEx models as 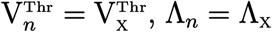, and 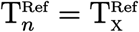, for all 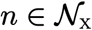 and x ∈ {e, i}. Note that all inhibitory neurons are considered to be non-adapting, that is, *η_n_* = 0 and *γ_n_* = 0 for all 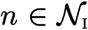. Similarly, for all synapses of type Y-to-x, where x, y ∈ {e, i}, that means for all 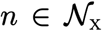 and 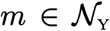, we set 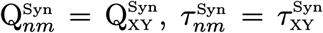, and 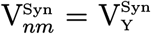. Note that, similar to the mean-field model, we assume that the synaptic reversal potentials do not depend on the type of the postsynaptic neurons.

We generate the elements of the internal network connectivity, 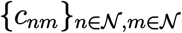, using the same connection probabilities used for the mean-field model. That is, we set *c_nm_* = 1 with probability P_xy_, as given in Table 1, when 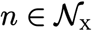 and 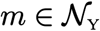, with x, y ∈ {e, i}. To set the elements of external connectivity, 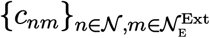, as well as the cardinality of 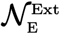, denoted by 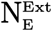, we first note that an in vivo network of size 10, 000 neurons can receive over 100, 000 afferent fibers from neurons outside of the network. The activities of the external neurons, however, are correlated. To lower the computational cost of simulating the network, and also to approximately take into account the correlation between external inputs, we assume that our spiking network receives input from only 1000, but independently spiking, input channels. That is, we set 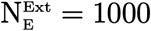. These input channels are then randomly connected to the neurons inside the network, with a connection probability of 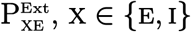. Therefore, *c_nm_* = 1 with probability 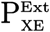 when 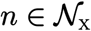 and 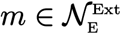, with x ∈ {e, i}.

We assume that spikes in each external input channel arrive randomly at Poisson-distributed time instances. To be able to compare the activity of the two models, we adjust the rate of spiking in each channel so that the average excitatory drive to neurons of the spiking network model becomes equal to the average background drive to the neurons of the mean-field model. For this, note that each neuron of type x in the mean-field model receives, on average, background inputs from 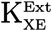 external excitatory neurons, each firing at an average rate of 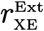 spikes per second. Correspondingly, each neuron in the spiking network receives background inputs from an average number of 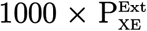 channels. If 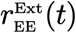 is set equal, or linearly proportional to 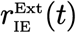 for all *t* ∈ [0, *T*], then we can choose the values for 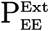 and 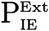 such that

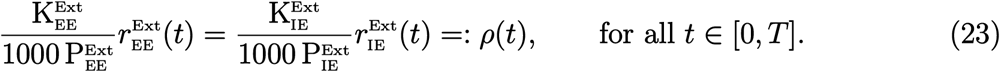

The scaling factors 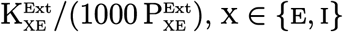, in (23) make the required adjustments on the external drive to the mean-field model so that *ρ*(*t*), as defined in (23), provides an equivalent drive to the spiking network. Therefore, we set the spike rate of each Poisson channel equal to *ρ*(*t*). For the numerical analysis presented in the Results section, we set 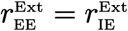 as in the mean-field model, and 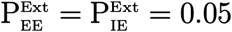 as given in Table 1.

## Results

The mean-field model (13)–(15) and the spiking network model (21)–(22), with biophysical parameter values given in Table 1, are used here to investigate how the overall balance of excitation and inhibition in cortical networks is affected by variations in some of the important physiological and structural factors of the network. We first show that the meanfield model with baseline parameter values is balanced with an activity rate typically observed in asynchronous irregular regimes, and is very responsive to changes in external inputs. We then perform the numerical analysis described in the Methods section and present how the network balance is affected by changes in key synaptic and structural parameters. We discuss the results and their biological implications in the Discussion section.

### The baseline balanced state

In order to investigate the effect of different synaptic and network parameters on the overall balance of excitation and inhibition, as described in next sections, it is important to first establish a reference balanced state for the network. Here, we demonstrate that the choice of baseline biophysical parameter values discussed in the Model Description section results in balanced state in the mean-field network activity, noting that our interpretation of the presence of overall balance in a network is based on observing the typical balanced network properties we described in the Introduction section. For this, we solve the mean-field equations (13)–(15) with the baseline parameter values given in Table 1. The transfer functions used in (13)–(15) are given by (11), with their arguments being computed using (1)–(10), (12), and the fit parameters given in Tables 2 and 3. We drive the network with excitatory background inputs of constant mean frequency 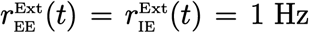 Hz, which presents a background activity at a level typically observed in irregularly spiking excitatory neurons at a cortical resting state [37].

Simulation of the mean-field model with the baseline setup described above identifies a stable equilibrium in the dynamics of the network, to which the mean-field activity of the network converges quickly. As we discussed in the Introduction section, computing the (mean) value of the important biophysical quantities of the network at this mean-field steady-state can provide a reasonably accurate estimate of the overall level of excitation and inhibition in the network. At this steady state, the excitatory neurons of the baseline model fire at the mean rate *p*_e_ = 1.15 Hz, and the inhibitory neurons fire at the mean rate *p*_i_ = 5.71 Hz. These rates of activity are close to the average excitatory firing rate of 1.09 Hz and inhibitory firing rate of 6.00 Hz obtained in a detailed simulation of the rat neocortical microcircuitry, when the constructed network is presenting balanced activity in an asynchronous irregular regime [37, Figure 17]; see also [29]. Moreover, our simulation of the spiking neuronal network that we construct equivalently to the mean-field model, as described in Model Description section, further confirms the presence of asynchronous irregular neuronal activity in the baseline model; see the rastergram shown in Figure 11a.

It is shown in the literature that excitatory and inhibitory synaptic conductances in the intact neocortex are well-balanced and proportional to each other [3]. Using (5) and (6), the mean value of excitatory and inhibitory synaptic conductances at the mean-field network equilibrium described above are calculated for the excitatory population as 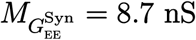 and 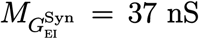, which are also equal to the values obtained for the inhibitory population. These mean conductance values are comparable to the experimentally measured values provided in [3]. They give an excitatory to inhibitory mean conductance ratio of 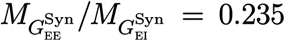, which is consistent with the experimental findings that imply inhibitory conductances are much larger than excitatory conductances [33, 53]. The results shown in [3], however, indicate a balanced conductance ratio of approximately equal to 1. The discrepancy between these results, as hinted in [3], may be because of the dominance of inhibitory synapses that can bias the results of experimental measurements. As we stated before in our discussion on the choice of parameter values for synaptic quantal conductances, the average number of inhibitory synapses per connection in our model is approximately four times larger than the average number of excitatory synapses per connection. If we assume this difference in the number of synapses proportionally biases experimentally measured conductances by a factor four, then the baseline ratio 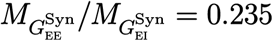 that we obtained in our simulation should be scaled by a factor of four to be comparable to the approximate ratio of 1 reported in [3]. We avoid such adjustment though, and in our analysis provided in the next sections we consider the ratio 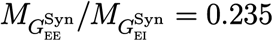 as a reference for the steady-state value of the balanced conductance ratio.

Experimental observations suggest that the dynamic balance of excitation and inhibition in local cortical networks keeps the neurons of the network in a depolarized state near their firing threshold, so that the network can be rapidly activated by external excitatory inputs and become involved in specific computational tasks [3, 38]. To ensure that the baseline balanced state in the mean-field model indeed corresponds to such a state of highly responsive network activity, we simulate the model with the same baseline parameter values as before, but with different values for the constant mean frequency of the excitatory inputs, 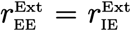. The resulting steady-sate values for different descriptive biophysical quantities of the model, obtained at the stable equilibrium of the equations for each input frequency value, are shown in Figure 1. First, it can be seen in Figure 1 that all biophysical quantities of the model, such as the mean firing rates, mean excitatory adaptation current, mean value and standard deviation of membrane potentials, and mean synaptic conductances take biologically reasonable values as the input frequency varies over a wide ranges of values. Second, the variation profile of the mean firing rates *p*_e_ and *p*_i_ shown in Figure 1a indicates that the overall activity of the neurons at the baseline background input frequency of 1 Hz, which is marked by dots in the graphs shown in Figure 1a, is indeed close to the firing threshold of the neurons. Last, relatively sharp changes in the mean firing activity of the neurons in response to different levels of excitatory input stimuli indicates that the baseline network is in a sufficiently responsive state. Moreover, although not shown here, our simulation results also verify that the mean-field dynamics of the network with baseline parameter values is sufficiently fast in responding to rapid fluctuations in the external inputs. A sample of the network firing response to fluctuations of approximate frequency 10 Hz is shown in Figure 10a. Such rapid response is also observed to fluctuations as fast as 20 Hz.

The mean inhibitory activity in the model rises in parallel to the mean excitatory activity as the mean frequency of external excitatory inputs increases to larger values beyond the background value. However, the results shown in Figure 1 imply that the neuronal interactions throughout the network control the overall level of inhibition in the network at a level that still allows for an elevated level of overall excitation—which is necessary for the network to be able to perform the processing task demanded by the external stimuli. This results in a change in the overall balance of excitation and inhibition toward higher excitation, as observed through the increase in the ratio between mean excitatory and inhibitory synaptic conductances, shown in Figure 1c. Nevertheless, the level of inhibition in the network remains sufficiently strong to prevent network instability and hyperactivity when the network receives an excessive amount of excitatory inputs from other cortical regions.

The observations made above confirm that our mean-field model with the baseline parameter values and background external inputs of mean frequency 1 Hz represents a network at a well-balanced state of overall excitation and inhibition. At this state, the network stays in an asynchronous irregular regime and is highly responsive to external cortical inputs, without undergoing internal instabilities when the level of external excitation increases. Therefore, we choose this state as the baseline balanced state of the mean-field model, and use it as a reference state to study how such a balanced state is disturbed by changes in the values of physiological and structural parameters of the network.

### Synaptic contributors to the balance of excitation and inhibition

The kinetics of synaptic activity in the conductance-based model we use here is governed by three main physiological factors, namely, synaptic decay time constants, synaptic quantal conductances, and synaptic reversal potentials. Variations in these physiological factors directly change the efficacy of synaptic communications between the neurons and hence have substantial impact on the dynamic balance of excitation and inhibition across the network. We investigate such impacts by showing how the steady-state balance between excitatory and inhibitory synaptic conductances—obtained at the stable equilibrium of the baseline meanfield activity described above—is affected by variations in each of these synaptic factors. In particular, we identify critical states of imbalanced activity which result in the loss of stability of the network equilibrium and lead to a transition of the network dynamics to an oscillatory regime.

The dynamics of the mean-field model (13)-(15) are highly dependent on the profiles of the transfer functions *F*_e_ and *F*_i_, which can change significantly if the physiological parameters of the synapses change. Hence, demonstrating the effects of variations in synaptic parameters on the profile of transfer functions (neuronal response curves) helps our understanding of how such variations affect the balance of mean-field activity in the network. Figure 2 illustrates how neuronal response curves vary with respect to changes in each of the three synaptic parameters we consider here. Variations in the decay time constants of inhibitory synapses, as shown in Figure 2a, changes the gain (sensitivity) of both inhibitory and excitatory neurons by changing the slope of their response curves. Additionally, such variations also change the excitability of the neurons by horizontally shifting their response curves. Similar effects are observed when the decay time constants of excitatory neurons change, however, with changes in neuronal excitability being less pronounced in this case. Figure 2b shows that changes in synaptic quantal conductances similarly affect gain and excitability of the neurons, with a higher sensitivity of the response curves being observed with respect to variations in excitatory quantal conductances. Changes in the synaptic reversal potentials, as shown in Figure 2c, significantly alter the gain of the neurons but have a lesser impact on their excitability. Changes in the gain of the neurons when the excitatory synaptic reversal potentials are increased or decreased from their baseline values are pronounced, and occur monotonically. However, gain changes with respect to variations in inhibitory reversal potentials appear to be non-monotonic. At lower output frequency values, both increasing 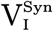 above its baseline value, and decreasing it below its baseline value, result in an increase in the gain of the neurons.

**Figure 2:**
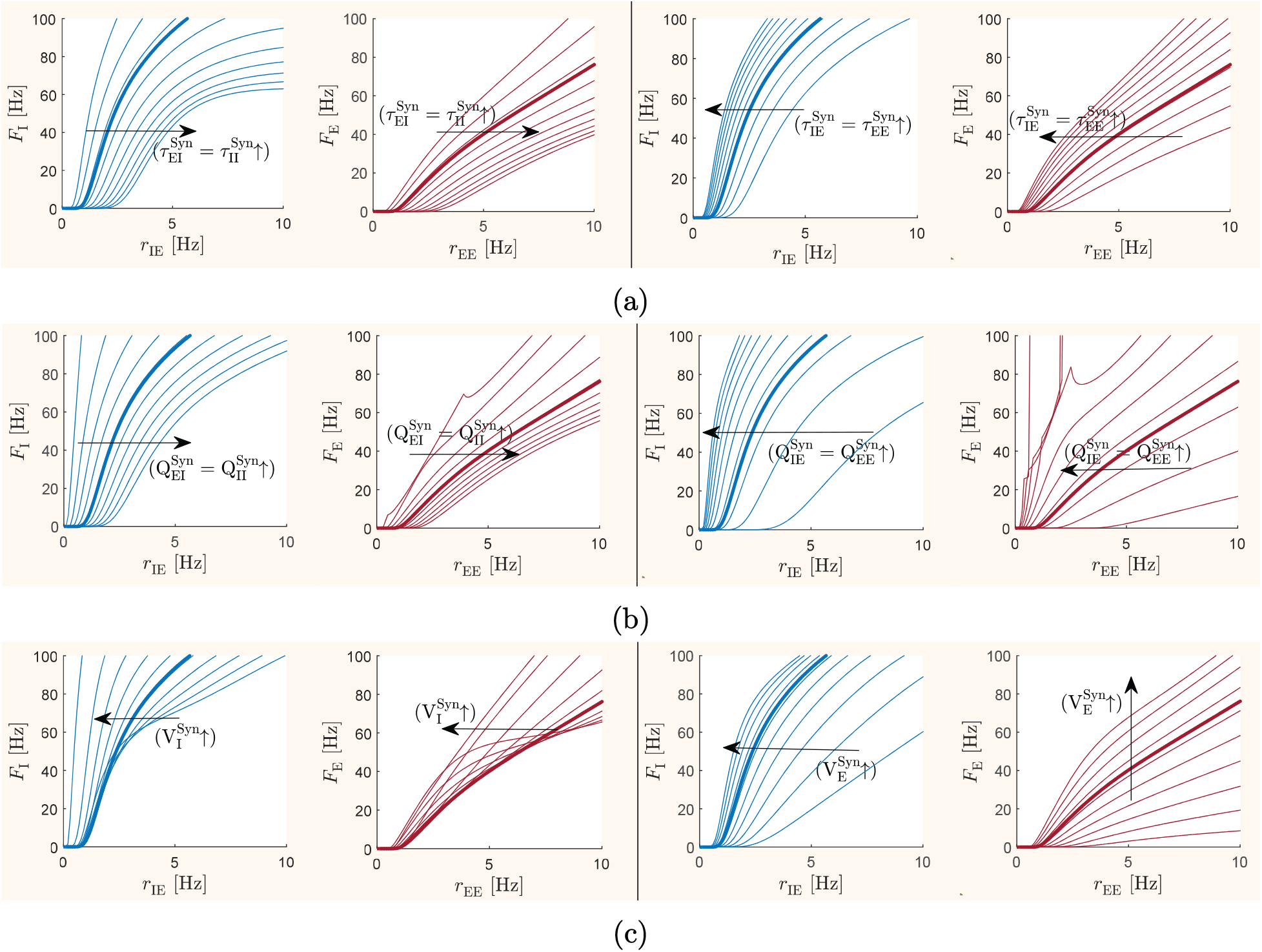
Variations in the response curves of neurons as a result of changes in synaptic parameters. For all graphs, the mean inhibitory spike rates received by both populations are fixed at the typical value *r*_ei_ = *r*_ii_ = 6 Hz. The mean excitatory spike rate received by each population is varied over a plausible range of values to obtain each neuronal response curves *F*_x_, *x* ∈ {e, i}, which are calculated using (11) and the fit parameters given in Table 3. All parameter values involved in the calculation of *F*_x_, except those specified on each graph, take their baseline values as given in Table 1. Thick curves in each graph show the response curves obtained at baseline synaptic parameter values. Other curves in each graph illustrate variations in the shape of the response curves as a synaptic parameter changes. Arrows indicate variations corresponding to 10 evenly distributed incremental changes in the parameter values specified in each graph. (a) Response curves with respect to variations in inhibitory (shown on the left side of the panel) and excitatory (shown on the right side of the panel) synaptic decay time constants. The value of 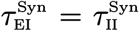 is varied from 5 ms to 18 ms, and the value of 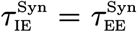 is varied from 1 ms to 3 ms. (b) Response curves with respect to variations in inhibitory and excitatory synaptic quantal conductances. The value of 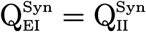 is varied from 1 nS to 25 nS, and the value of 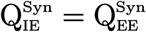 is varied from 1 nS to 8 nS. (c) Response curves with respect to variations in inhibitory and excitatory synaptic reversal potentials. The value of 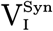 is varied from −103 mV to −53 mV, and the value of 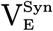 is varied from −30 mV to 20 mV.

Local inhibitory sub-networks are known to play a key role in stabilizing the dynamics of local networks and coordinating the flow of activity across cortical areas [8,12,28,34,36,54]. Therefore, in what follows, we initially perform our analysis based on codimension-one continuation of the baseline equilibrium state with respect to variations in each of the inhibitory synaptic parameters. Then, we extend the analysis to codimension-two by additionally considering variations in excitatory synaptic parameters.

#### Effect of synaptic decay time constants

The decay time constant of a synapse determines how long the activity initiated in the synapse by an incoming action potential (spike) will last. Hence, the decay time constant has a significant impact on the efficacy of the synapse, which can also be directly implied from the mean synaptic conductance equations (5) and (6). Moreover, changes in synaptic decay time constants also change the standard deviation and autocorrelation time constants of membrane potential fluctuations, as implied from equations (9) and (10). As a result, the transfer functions (response curves) of neurons are highly impacted by variations in the decay time constants, which is confirmed by the results shown in Figure 2a. To show how the decay time constants of the synapses then impact the global dynamics of the network, and how they contribute to maintaining or disturbing the overall balance of excitation and inhibition, we first continue the stable equilibrium of the baseline balanced network with respect to variations in inhibitory decay time constants 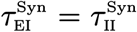. The results are shown in Figure 3.

**Figure 3:**
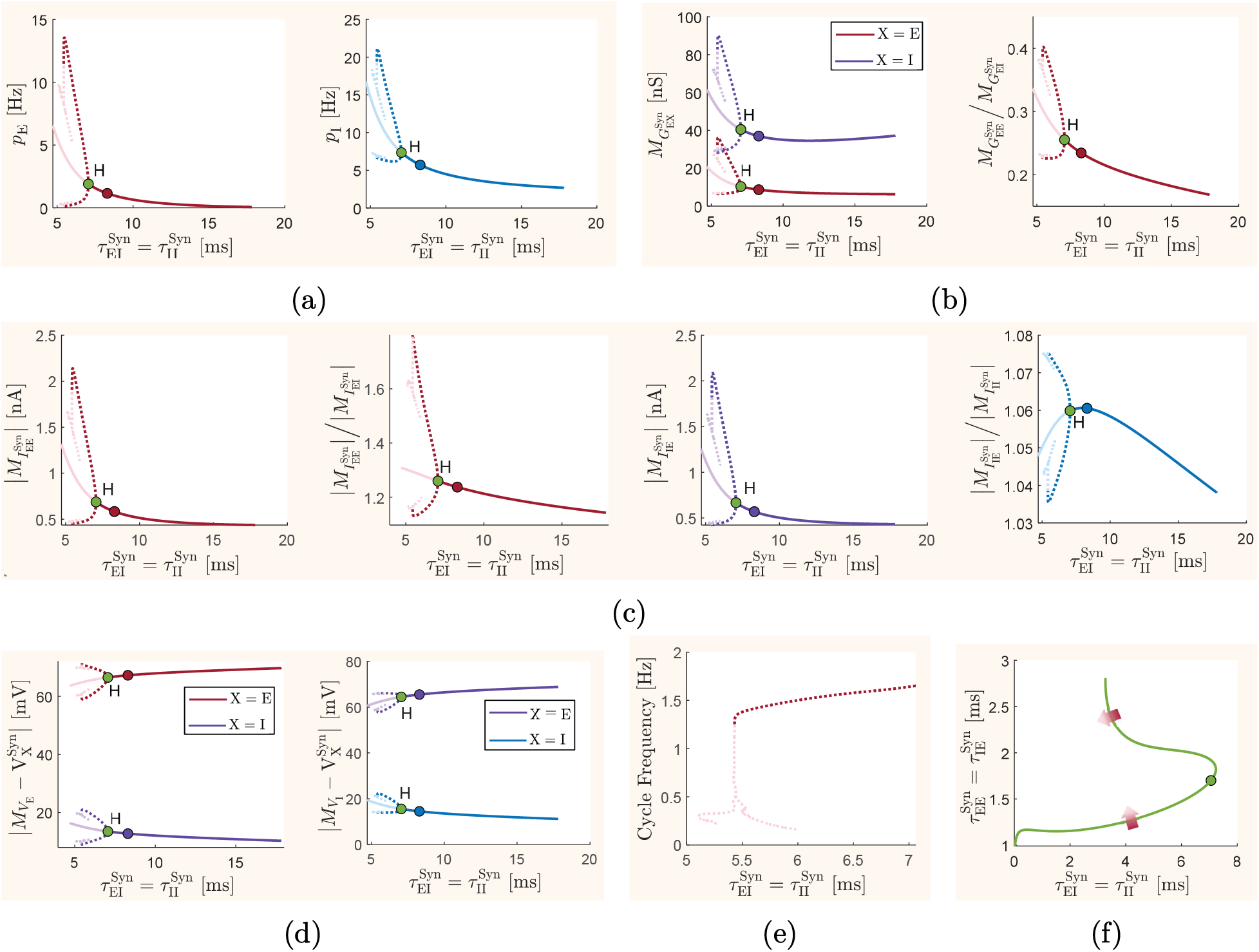
Effect of variations in synaptic decay time constants on the long-term meanfield activity of the network. All parameter values of the mean-field model, except for the synaptic decay time constants that are explicitly specified in each graph, are set to their baseline values given in Table 1. The model is driven by background inputs of constant mean frequency 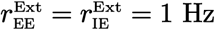. In graphs (a)–(e), the decay time constants of inhibitory synapses made on neurons of both excitatory and inhibitory populations are set to take the same value, which is varied as the bifurcation parameter. The resulting codimension-one continuation of the network equilibrium and emerging limit cycles are shown for different quantities. Curves of equilibria are shown by solid lines, and the minimum and maximum values that each quantity takes on the limit cycles are shown by dotted lines. Dark-colored segments of each curve indicate stable equilibria/limit cycles, whereas light-colored segments indicate unstable equilibria/limit cycles. Unlabeled dots in each graph correspond to the baseline parameter values 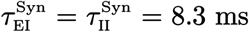. Points labeled by H are Hopf bifurcation points. We show bifurcation diagrams for (a) mean excitatory firing rate *p*_e_ and mean inhibitory firing rate *p*_i_, (b) mean synaptic conductances 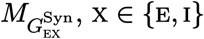, of the excitatory population, and the ratio between the two conductances, (c) absolute value of the mean excitatory synaptic currents, 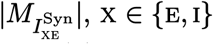, as well as the ratio between mean excitatory and inhibitory synaptic currents, and (d) mean excitatory and inhibitory synaptic driving forces 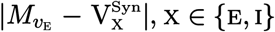, in the excitatory population, as well as such driving forces 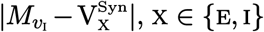, in the inhibitory population. (e) Frequency of the limit cycles originating from the Hopf bifurcation point. (f) Codimension-two continuation of the Hopf bifurcation point when the excitatory synaptic decay time constants are also allowed to vary as a bifurcation parameter. The point marked by a dot has the baseline excitatory parameter value of 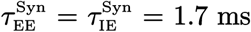. Arrows indicate transition from stable (dark) to unstable (light) equilibria as the parameters are varied across the curve.

The response curves given in Figure 2a show that increasing the value of 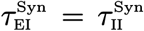 decreases the gain and excitability of inhibitory and excitatory neurons. As a result, the mean firing rates of both excitatory and inhibitory populations decrease, concurrently, with increasing 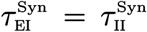, as shown in Figure 3a. This paired variation in the mean firing activity of the populations, however, does not necessarily imply that the overall balance of excitation and inhibition remains unchanged with respect to changes in inhibitory decay time constants. The ratio of the mean excitatory to mean inhibitory synaptic conductances in the excitatory population, 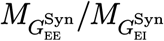, shown in Figure 3b, implies that the excitationinhibition balance moves toward over-inhibition as the efficacy of inhibitory synapses grows with increasing 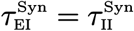. Indeed, unlike the profile of the inhibitory firing rate *p*_i_, at large values of 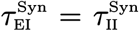 the mean inhibitory synaptic conductance 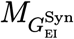 shown in Figure 3b rises slightly as 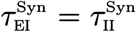 increases. It should be noted that mean synaptic conductances in the inhibitory population, 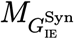 and 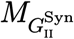, which are not shown in Figure 2a, take the same values as those in the excitatory population due to the same choices of connection probabilities in our model.

On the opposite direction, decreasing the value of inhibitory synaptic decay time constants increases the ratio 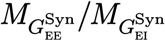 and moves the excitation-inhibition balance toward more excitation. The excessive excitation in the network then drives the network’s dynamics to a critical state, at which further reduction in 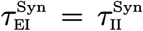 results in an overall level of excitation that cannot be effectively balanced by the increased level of inhibition. At this critical state, the network’s dynamics undergoes a phase transition to an oscillatory state, identified by a Hopf bifurcation point in the codimension-one continuations shown in Figure 3. The limit cycle originated from the Hopf bifurcation point has also been continued, and the curves of minimum and maximum values taken by different biophysical quantities of the model on the resulting cycles are shown in Figure 3. When the inhibitory synaptic decay time constants decrease below the critical value 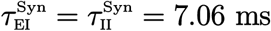, which is the value at which the Hopf bifurcation occurs, the previously stable equilibrium of the model becomes unstable. As a result, the orbits of the system depart the vicinity of the equilibrium and converge to a stable limit cycle on the curves of cycles originated from the Hopf bifurcation. Figure 3e shows that the stable oscillations on this cycle lie in the delta frequency band (1 - 4 Hz). It should be noted that further continuation of the curves of limit cycles, as partially shown in Figure 3, detects a fold bifurcation of limit cycles and emergence of unstable limit cycles at low values of 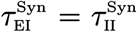. However, analyzing the dynamics of the model at such values is not pertinent to the purpose of this paper. In fact at such values the mean-field model loses its validity in accurately predicting the emerging dynamic regimes, since the quantity under the square root in (9) approaches negative values.

The results shown in Figure 3d imply that the mean membrane potential of both excitatory and inhibitory populations decreases with increases in the value of 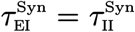. This means that, on average, neurons in the network become more hyperpolarized when the level of inhibition in the network is enhanced by increasing the efficacy of the inhibitory synapses. Since in this study the values of synaptic reversal potentials are kept fixed, the reduction in the mean membrane potential of the neurons due to an increase in 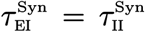 yields a reduction in the mean electrochemical driving force of the inhibitory synapses in both populations, 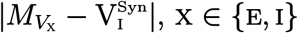, and a rise in the mean driving force of the excitatory synapses, 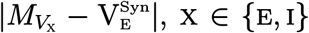. The curves of the equilibrium driving forces shown in Figure 3d illustrate such profiles of variations.

The absolute values of the mean synaptic currents at the equilibrium, calculated using (16) and (17), are shown in Figure 3c. As discussed above and illustrated in Figures 3b and 3d, the steady-state values of the two contributing components of these currents, the mean synaptic conductances and the mean synaptic driving forces, change in opposite directions with increases in 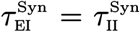. The combination of these two components, however, results in steady-state mean synaptic currents which decrease in absolute value as 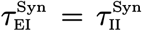 is increased, as we can see in the curves of equilibria shown in Figure 3c; note that inhibitory currents are not shown separately in Figure 3c as their profile is similar to that of excitatory currents. In particular, despite the increase in mean driving forces of the excitatory synapses, the mean excitatory synaptic currents decrease as 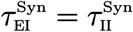 is increased. This implies that the effect of decreasing excitatory conductances, shown in Figure 3b, dominates the effect of increasing mean excitatory driving forces, hence resulting in the net decrease of excitatory synaptic currents.

Figure 3c also shows that the steady-state ratio between the absolute values of mean excitatory and mean inhibitory synaptic currents in the inhibitory population, 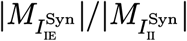, does not significantly change with respect to 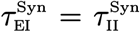, whereas the ratio between these currents in the excitatory population, 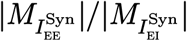, decreases with increasing 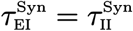. Therefore, similar to the ratio 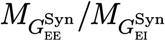 between synaptic conductances, here the ratio 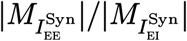 between synaptic currents also indicates shifts in the level of excitation-inhibition balance, toward over-inhibition or over-excitation.

The results provided above describe only the effect of inhibitory synaptic decay time constants on the network balance. Nevertheless, the excitation-inhibition balance in the network can also be affected by changes in the excitatory synaptic decay time constants. It is implied from the the neuronal response curves shown in Figure 2a that changes in excitatory and inhibitory decay time constants have opposite impacts on the gain of neurons. Therefore, with fixed values of 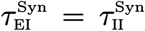, the mean firing rates of both inhibitory and excitatory populations increase concurrently with increases in 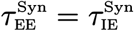. However, similar to changes in inhibitory decay time constants described above, it is not intuitively possible to accurately predict whether the network balance will be maintained during such paired changes in the mean firing rates of the populations; and if not, to which direction, either over-inhibition or over-excitation, the balance will shift. In fact, the relative changes between the magnitude of the mean excitatory and inhibitory response curves, and the varying sensitivity of these curves to changes in synaptic decay time constants—which results from complicated interactions in the network between adaptive and non-adaptive dynamic neurons—can result in non-intuitive changes in the network balance.

To demonstrate the joint effect of both excitatory and inhibitory synaptic decay time constants on network balance, we perform a codimension-two continuation of the Hopf bifurcation point that was detected in the codimension-one bifurcation analysis described before. We consider the same values for the decay time constants of excitatory synapses in both populations, 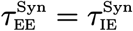, and vary it as a second bifurcation parameter. The resulting curve of Hopf points is shown in Figure 3f.

It is observed that, when 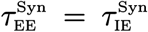 is reduced from its baseline value, the critical transition point (Hopf bifurcation) in the dynamics of the network occurs at lower values of 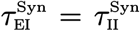. This implies that lower values of excitatory synaptic decay time constants reduce the overall level of excitation in the network, so that this reduction in excitation effectively compensates for a rise in excitation caused by decreases in inhibitory decay time constants 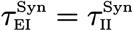. Therefore, due to this compensatory effect, lower values of excitatory synaptic decay time constants allow for the network to maintain a stable balanced state at much lower values of inhibitory decay time constants.

The upper part of the curve of Hopf points in Figure 3f shows a less intuitive dynamics for larger values of 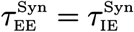 than its (approximately) baseline value. Although not shown here, codimension-one continuations of the baseline equilibrium with respect to variations in 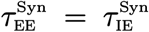, but with fixed values of 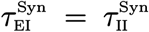, still show an increase in the level of excitation at higher values of 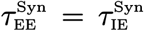 and a shift in the balance of excitation and inhibition to hyper-excitation. However, as it can be seen in Figure 3f, at large values of inhibitory synaptic decay time constants, including the baseline value 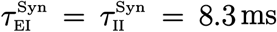, this shift to hyper-excitation does not drive the network’s dynamics to an oscillatory regime. Both an oscillatory state and a non-oscillatory hyper-excited state can be observed in the network activity when 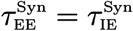 is increased from small values to large values while 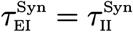 is at about 4 – 7 ms. It should be noted, however, that the level of stable hyperexcitation observed at large values of excitatory synaptic decay time constants corresponds to conductance ratios 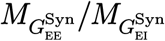 that may not be biologically plausible.

Importantly, the curve of Hopf bifurcation points in Figure 3f implies that the baseline parameter values 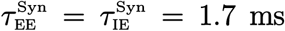 and 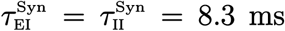 are critical values for network stability, in the sense that the network remains quite responsive at these values, while relatively small reduction in the mean decay time constant of inhibitory neurons transitions the network’s dynamics to an oscillatory state.

#### Effect of synaptic quantal conductances

The quantal conductance of a synapse is the peak membrane conductance change in a neuron caused by receiving a single spike from a presynaptic neuron. Therefore, changes in the quantal conductance of a synapse directly modulate the strength of the synapse. This is also implied from the mean synaptic conductance equations (5) and (6). It should be noted that, as described in the Model Description section, the quantal conductances Q_xy_, x, y ∈ {e, i}, that are incorporated in the mean-field model we use here are in fact quantal conductances per connection, that means, they are equal to the quantal conductances per synapse times the number of synapses per connection. As a result, Q_xy_, x, y ∈ {e, i}, in our study can be altered both by changes in the number of synapses and by modulations of the synaptic peak conductances.

The mean synaptic conductances (5) and (6) change with synaptic quantal conductances in the same way as they change with synaptic decay time constants. However, standard deviation and autocorrelation time constants of membrane potential fluctuations change differently with synaptic quantal conductances. In general, similar to the effect of increases in 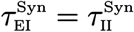, the response curves given in Figure 2b show that the gain and excitability of both inhibitory and excitatory neurons decrease with increasing 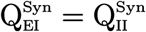. Therefore, we expect that the effects on network balance caused by varying synaptic quantal conductances to be similar to the effects caused by changing synaptic decay time constants. This can be seen in Figure 4, which shows bifurcation diagrams of key biophysical quantities with inhibitory quantal conductances 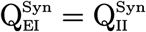 varied as the bifurcation parameter.

**Figure 4:**
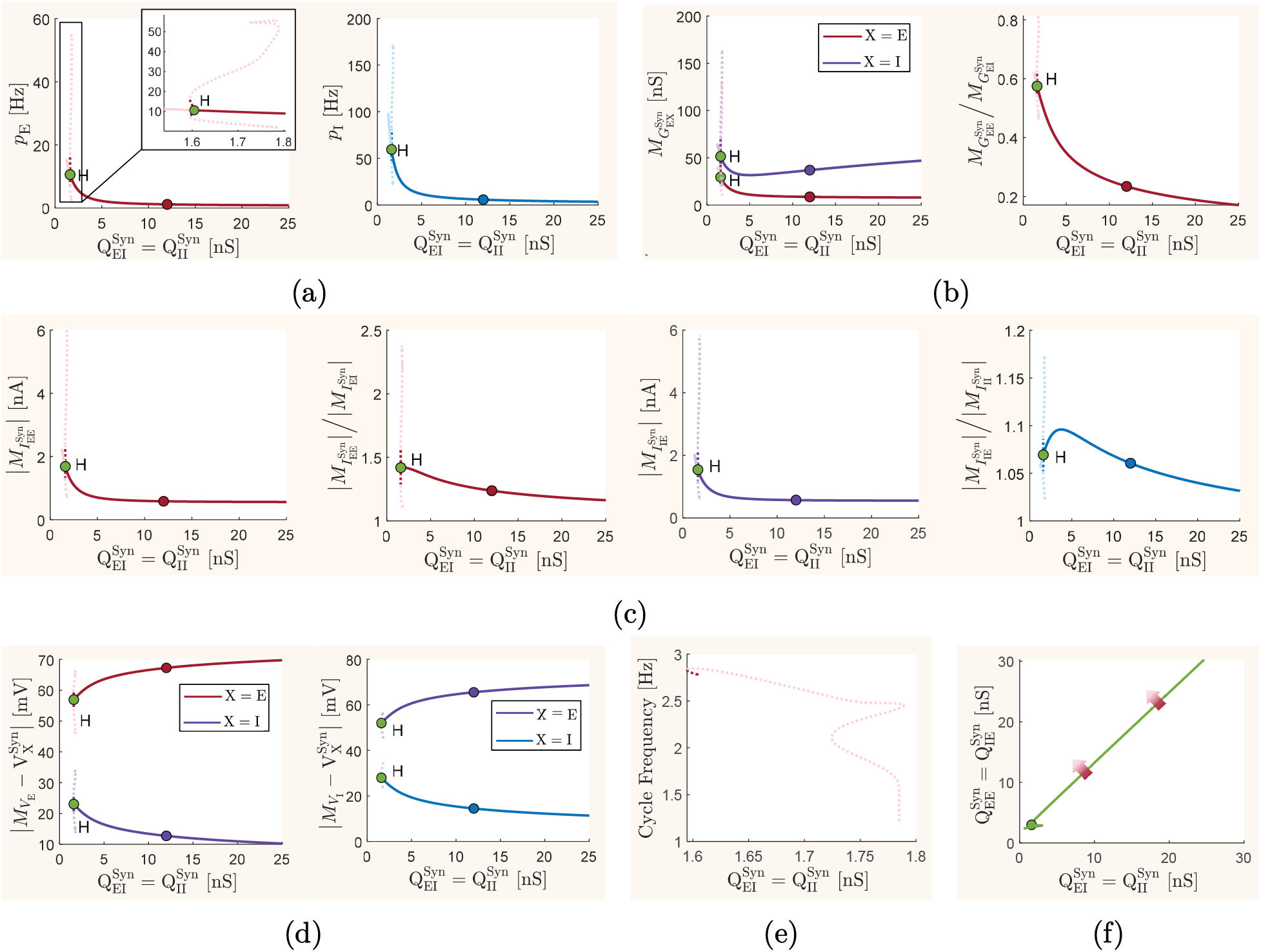
Effect of variations in synaptic quantal conductances on the long-term mean-field activity of the network. All parameter values of the mean-field model, except for the synaptic quantal conductances that are explicitly specified in each graph, are set to their baseline values given in Table 1. The model is driven by background inputs of constant mean frequency 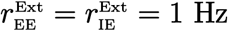. The same description as given in Figure 3 holds for the graphs presented here, with the only difference being that here the bifurcation parameter for codimension-one continuation is the inhibitory quantal conductances 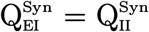, with the baseline value of 12 nS, and the additional bifurcation parameter for codimension-two continuation is the excitatory quantal conductances 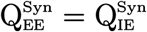, with the baseline value of 3 nS.

In a rather similar way to what we observed in Figure 3 when 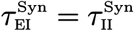 was changed, we also see here that when the value of inhibitory quantal conductances 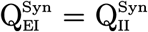 increases the following changes occur in the steady-state values of different quantities of the model: the mean firing rates of both excitatory and inhibitory populations decrease concurrently; the ratio of mean excitatory to mean inhibitory synaptic conductances decreases; the absolute value of mean synaptic currents decreases; the ratio of the absolute value of mean excitatory to the absolute value of mean inhibitory synaptic currents in the excitatory population decreases; the mean electrochemical driving force of the inhibitory synapses in both populations decreases; and the mean driving force of the excitatory synapses increases. As a result, the overall excitation-inhibition balance in the network shifts toward over-inhibition as the inhibitory synapses are strengthened by increasing 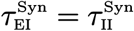.

In contrast to the high sensitivity of the network’s stability to reductions in decay time constants of the inhibitory neurons, the bifurcation diagrams of Figure 4 imply that the stability of the network’s equilibrium is relatively robust to decreases in the value of inhibitory quantal conductances. Comparing the curves of equilibrium values for the ratios 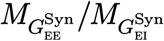 and 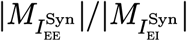 shown here in Figures 4b and 4c, respectively, with those shown in Figures 3b and 3c, we can see that significantly higher level of overall excitation can be maintained in the network at low values of 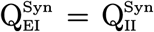 before a transition in the network’s dynamics to an oscillatory regime occurs. However, the loss of stability of the network’s equilibrium at very low values of 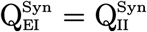 appears to be more critical. Stable delta-band oscillations can exists only for very short range of values of 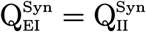.

The neuronal response curves given in Figure 2b show that changes in excitatory and inhibitory synaptic quantal conductances have opposite impacts on the gain and excitability of the neurons. Unlike the non-intuitive dynamics that we observed in Figure 3f when decay time constants of the synapses were jointly changed, the curve of Hopf bifurcation points shown in Figure 4f, when both 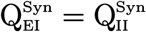 and 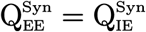 are free to change, shows an intuitively expected dynamics. Increasing the value of excitatory quantal conductances results in a rise in the level of excitation in the network, which then requires stronger inhibitory synapses for maintaining the network stability and avoiding transitions to oscillatory states. As a result, the Hopf bifurcation point occurs at higher values of 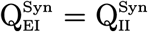 when 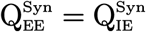 is increased. Moreover, with any fixed values of inhibitory quantal conductances, a phase transition occurs in the network’s dynamics if the quantal conductances of excitatory synapses increase beyond a Hopf bifurcation value. However, the baseline parameter values 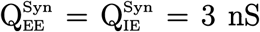 and 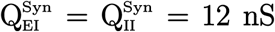 appear to be significantly below the curve of Hopf bifurcation points in Figure 3f, implying that the network stability at the baseline state is sufficiently robust to changes in both inhibitory and excitatory synaptic quantal conductances.

#### Effect of synaptic reversal potentials

Changes in reversal potentials of synapses modulate the electrochemical driving force of the synapses and also directly affect the mean membrane potential of the neurons across the network, as implied from (8). It can also be seen through (9) and (10) that changes in synaptic reversal potentials also affect the standard deviation and autocorrelation time constants of neuronal membrane potential fluctuations. Therefore, the characteristic response curves of the neurons and the dynamics of overall excitation-inhibition balance in the network can be significantly affected by changes in the reversal potential of the synapses. To show such impacts on the global dynamics of the network, we perform bifurcation analysis with respect to variations in synaptic reversal potentials, as we have performed with respect to variations in the other synaptic parameters. That is, we first continue the baseline stable equilibrium of the network with respect to variations in the value of inhibitory reversal potential 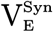, and then extend the results by performing a codimension-two continuation of the detected Hopf bifurcation points by additionally allowing variation of the excitatory reversal potential 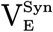. The results are shown in Figure 5.

**Figure 5:**
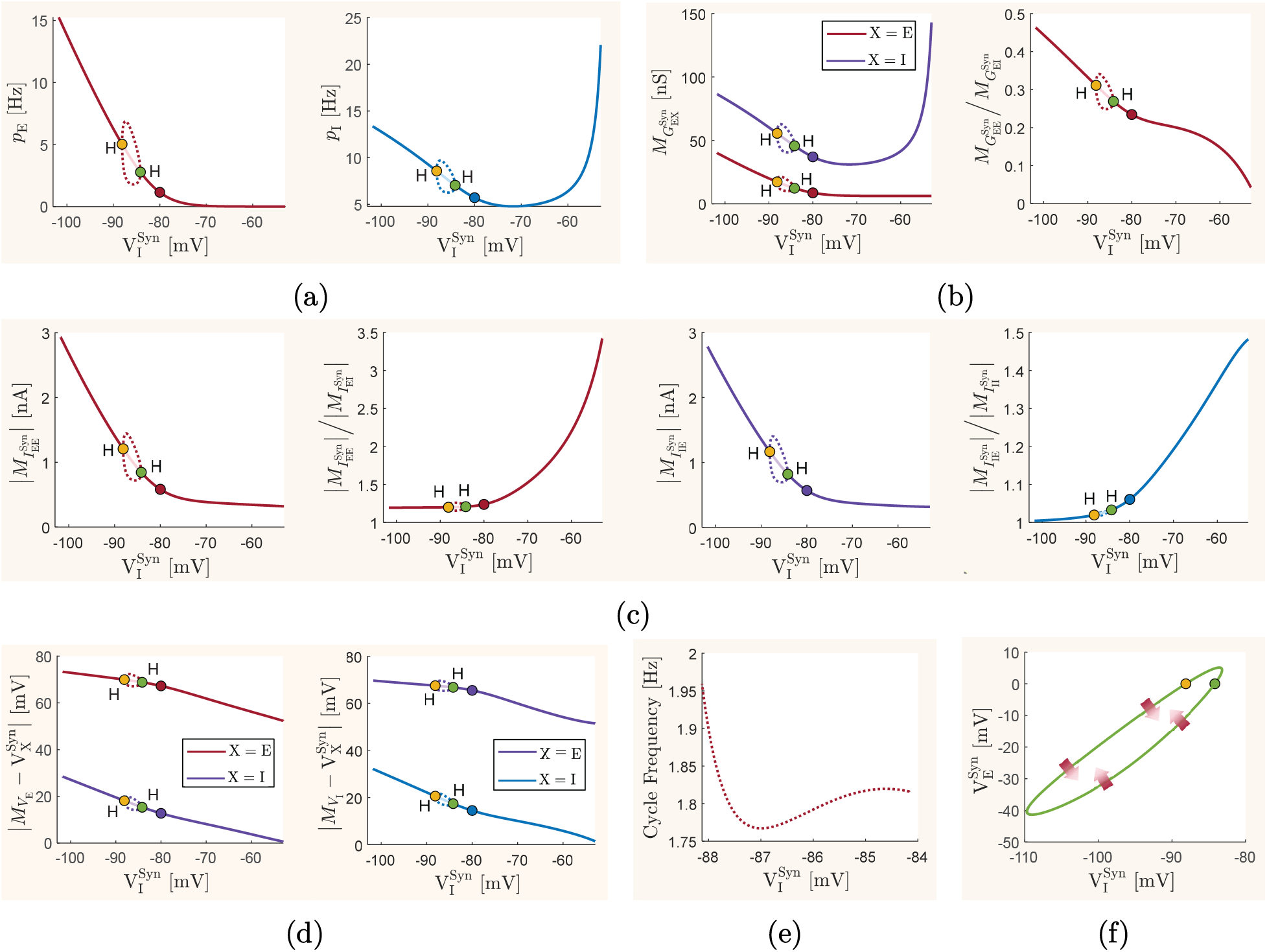
Effect of variations in synaptic reversal potentials on the long-term mean-field activity of the network. All parameter values of the mean-field model, except for the synaptic reversal potentials that are explicitly specified in each graph, are set to their baseline values given in Table 1. The model is driven by background inputs of constant mean frequency 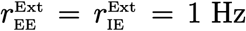. The same description as given in Figure 3 holds for the graphs presented here, with the only difference being that here the bifurcation parameter for codimension-one continuation is the inhibitory reversal potential 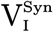, with the baseline value of −80 mV, and the additional bifurcation parameter for codimension-two continuation is the excitatory reversal potential 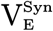, with the baseline value of 0 mV.

The neuronal response curves in Figure 2 show that, within a wide range of plausible firing rate frequencies, the gain of inhibitory and excitatory neurons increases with both increasing and decreasing changes in the value of 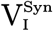 from its baseline value, although the increase in the gain of inhibitory neurons caused by a reduction in 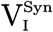 is less pronounced. This may suggest that the mean firing rate of the inhibitory and excitatory populations will increase with 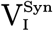 changes in either directions. However, the curves of steady-state mean firing rates shown in Figure 5a do not confirm this intuitive prediction. Although the steady-state value of *p*_i_ increases when 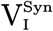 decreases below its baseline value, as well as when 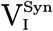 increases sufficiently above its baseline value, the steady-state value of *p*_e_ is strictly decreasing with increasing 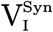. In particular, these profiles of variations in *p*_e_ and *p*_i_ have counter-intuitive relations with the steady-state values of the mean membrane potentials. Since 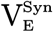 is fixed at the constant value of 0 mV in the codimension-one analysis presented here, the curves of excitatory driving forces shown in Figure 5d imply that, on average, both excitatory and inhibitory neuronal populations monotonically become more depolarized with increases in 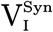, which is a pattern of variation not directly implied from the mean firing rate of the neurons.

An explanation for the network interactions that could result in the steady-state curves described above can be as follows. Although the neurons become further depolarized with increases in 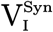, the driving forces of inhibitory synapses, 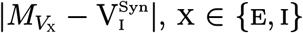, shown in Figure 5d keeps decreasing to very small values as 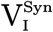 increases to large values. This results in significant reduction in inhibitory synaptic currents at large values of 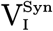, despite the rise in the firing rates of the inhibitory neurons due to their increased gain. As a result, a stable balance of excitatory and inhibitory synaptic currents can be achieved if the mean firing rate of excitatory neurons significantly reduces to near zero, as shown in Figure 5a. It should be noted, however, that in the studies presented here a constant level of excitation is always provided to all neurons of the network by the excitatory background inputs of constant frequency 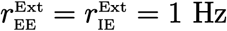. In the opposite direction, when 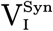 decreases to low values, the mean driving force of inhibitory synapses increases to sufficiently large values, which allow for inhibitory synaptic currents that are large enough to be balanced by larger excitatory synaptic currents. Therefore, a stable balance of currents can be achieved at low values of 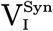, with both *p*_e_ and *p*_i_ taking relatively large values due to the increased neuronal gains. This increase in the mean firing rates of the neurons with decreases in 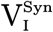 to sufficiently low values can be seen in Figure 5a.

The steady-state value of the ratio 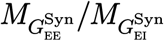 between mean excitatory and inhibitory conductances, as shown in Figure 5b, decreases with increasing 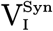. Consistent with the explanation provided above, this implies that the balance of excitation and inhibition in the network moves monotonically with 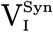, to over-inhibition for large values of 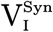, and to over-excitation for small values of 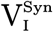. However, unlike what we observed in Figures 3c and 4c for changes in synaptic decay time constants and quantal conductances, here the monotonic shift in the level of excitation-inhibition balance cannot be correctly implied from the ratio 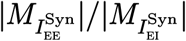 between synaptic currents in the excitatory population, as shown in Figure 5c.

The bifurcation diagrams of Figure 5 further reveal the presence of two Hopf bifurcation points. When 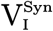 decreases below the Hopf point denoted by green dots in Figure 5, the network’s dynamics transitions to a stable oscillatory behavior. Figure 5e shows that the frequency of these oscillations are in the delta band. However, the emergent delta oscillations here are of significantly lower magnitude than those arising from changes in inhibitory synaptic decay time constants, as shown in Figure 3. Moreover, the curve of stable limit cycles calculated here vanishes at a second Hopf point when 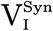 decreases further. This implies that reductions in the value of inhibitory synaptic reversal potentials is less critical to the stability of the network. In fact, a stable yet high level of excitation can be present in the network at low values of 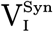.

Finally, codimension-two continuation of the two Hopf bifurcation points detected in the analysis described above implies that a qualitatively similar network dynamics is observed at a wide range of values of excitatory synaptic reversal potentials. This is shown in Figure 5f. Unlike changes with respect to 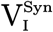, the neuronal response curves in Figure 2c show monotonic changes in the gain of neurons with respect to 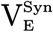. When 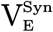 decreases, the gain of neurons decreases significantly. The resulting reduction in the level of excitation then allows for further reduction in 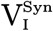 before the network transitions to an oscillatory regime. Therefore, the two Hopf bifurcation points occur at lower values of 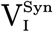 when 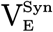 decreases.

### Structural contributors to the balance of excitation and inhibition

The mean-field model we use in our study represents a network of randomly connected neurons with relatively sparse connectivity. The dynamics of such network can be significantly affected by the structural factors that determine the overall topology of the network. These factors include the relative number of excitatory and inhibitory neurons in the network and the density of connectivity between the neurons. By employing bifurcation analysis similar to our previous investigations on the effect of physiological parameters intrinsic to synapses and neurons, we analyze how the steady-state balance of excitation and inhibition obtained at the baseline network structure is altered by changes in each of the structural factors determining the overall level of recurrent neuronal interactions in the network.

#### Effect of the ratio between the number of inhibitory and excitatory neurons

Knowing the crucial role of inhibitory neurons in stabilizing the excitatory interactions across the network, the ratio between the total number of inhibitory and excitatory neurons is expected to have significant impacts on the excitation-inhibition balance. Here, with a fixed number of neurons N = 10000, we change the inhibitory proportion N_i_/N of the total network population and observe how the steady-state mean-field activities in the network change accordingly. The resulting codimension-one bifurcation diagrams for different quantities of the model are shown in Figure 6.

**Figure 6:**
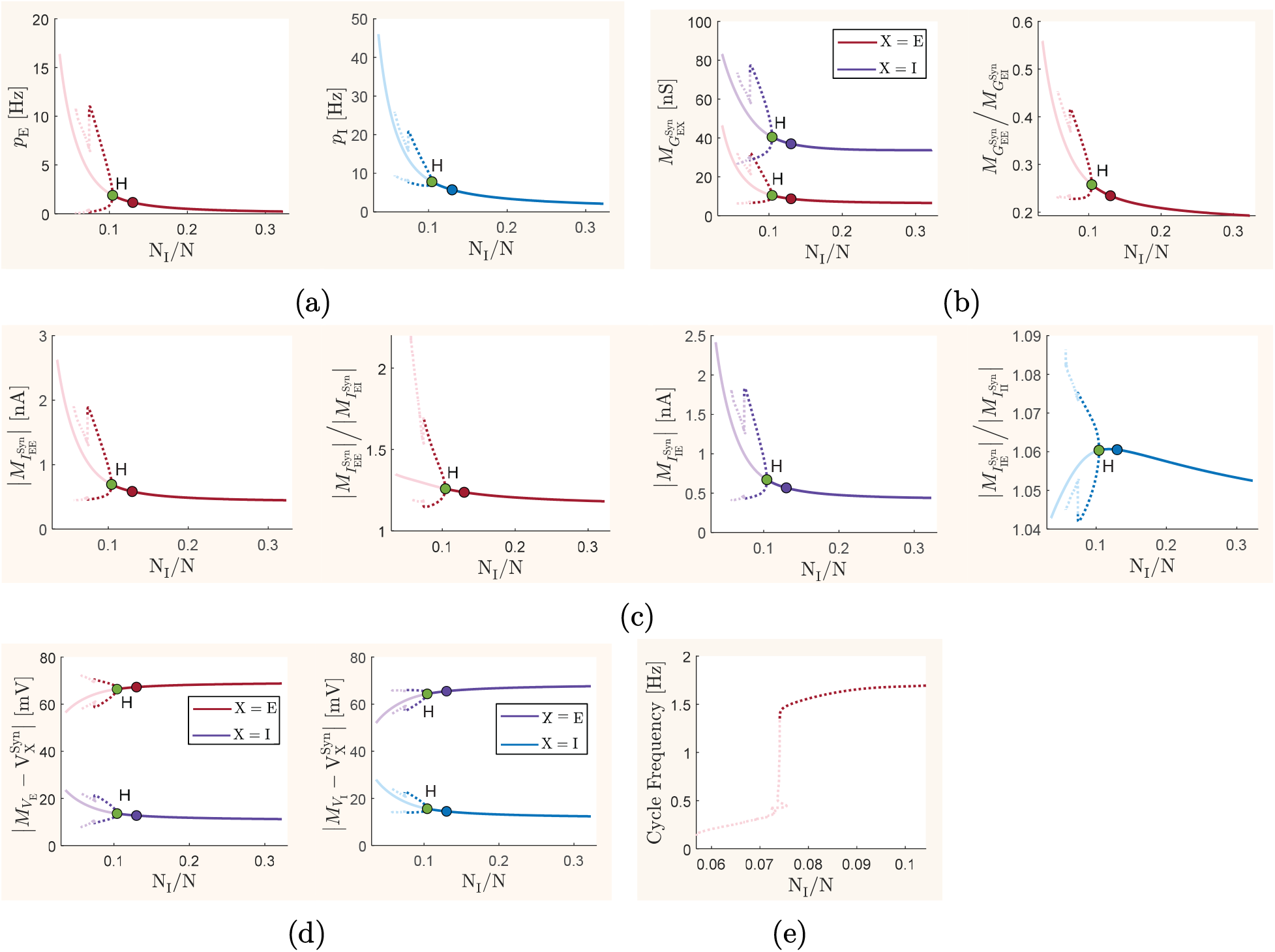
Effect of variations in the ratio between the number of inhibitory and excitatory neurons on the long-term mean-field activity of the network. All parameter values of the mean-field model, except for the number of inhibitory and excitatory neurons, are set to their baseline values given in Table 1. The total number of neurons is fixed at N = 10000. The inhibitory proportion of neurons in the network, N_i_/N, is varied over a reasonable range of values. The model is driven by background inputs of constant mean frequency 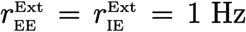. The same description as given for graphs (a)–(e) in Figure 3 holds for the graphs presented here, with the only difference being that here the bifurcation parameter for codimension-one continuation is N_i_/N, with the baseline value of 0.13.

Since N is fixed, a change in the ratio N_i_/N alters both N_e_ and N_i_. Therefore, considering (1) and (2), the mean values of both excitatory and inhibitory synaptic conductances given by (5) and (6), respectively, change with N_i_/N. Standard deviation and autocorrelation time constant of membrane potential fluctuations in both excitatory and inhibitory populations are also directly affected by changes in N_i_/N, as implied by (9), (10), (1), and (2). Moreover, a change in N_e_ and N_i_ directly modifies the dynamics of the covariance of firing rates, given by (14). Despite these different contributions in the equations of the model in comparison with those of the inhibitory synaptic decay time constants, the bifurcation diagrams obtained by changing N_i_/N closely resemble the diagrams obtained by changing 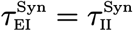. This can be seen by comparing Figure 6 with Figure 3. Therefore, when N_i_/N changes, the longterm dynamics of the network changes similarly to what we observed before as a result of changes in 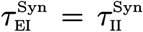. In particular, increasing the inhibitory proportion of the total network population N_i_/N from its baseline value moves the excitation-inhibition balance toward over-inhibition. Moreover, the stability of network activity appears to be critically sensitive to decreases in N_i_/N, so that relatively small reduction in N_i_/N causes a transition in network activity to stable oscillations in the delta frequency band.

The contribution of inhibitory neurons in regulating network activity is not only influenced by their relative population size, but also by their cellular properties and efficacy of the synapses that they make on other neurons, as we studied separately above. To see some of the joint effects of such structural and physiological factors on the stability of balanced activity in the network, we perform codimension-two continuation of the Hopf point detected in Figure 6 by additionally varying each of the inhibitory synaptic parameters as a second bifurcation parameter. The resulting curves of Hopf bifurcation points are shown in Figure 7.

**Figure 7:**
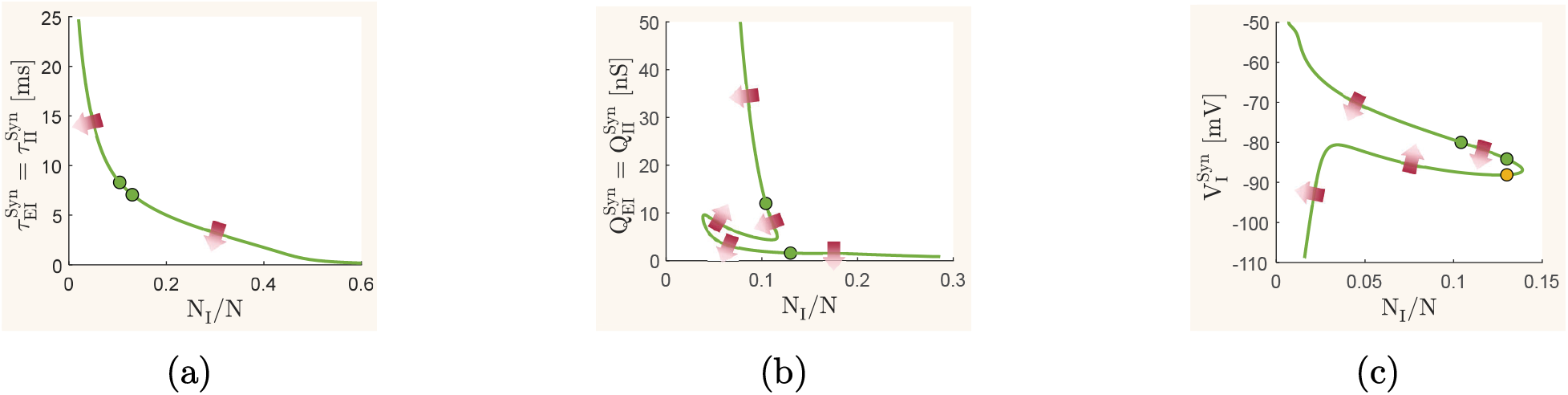
Joint effect of the synaptic parameters and the ratio between the number of inhibitory and excitatory neurons on the long-term mean-field activity of the network. Codimension-two continuation of Hopf bifurcation points are shown in each graph. The inhibitory proportion of neurons in the network, N_i_/N, is considered as one of the bifurcation parameters in all graphs. Different synaptic parameters are then chosen in each graph as the second bifurcation parameter. Arrows indicate transition from stable (dark) to unstable (light) equilibria as the parameters are varied across the curve. The points marked by dots are the same Hopf bifurcation points marked in Figures 3, 4, 5 and 6. (a) Curve of Hopf bifurcation points with decay time constant of inhibitory synapses chosen as the second bifurcation parameter. (b) Curve of Hopf bifurcation points with quantal conductance of inhibitory synapses chosen as the second bifurcation parameter. (c) Curve of Hopf bifurcation points with reversal potential of inhibitory synapses chosen as the second bifurcation parameter.

The results discussed before showed that changes in 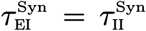 and N_i_/N have very similar impacts on the network’s dynamics. A decrease in 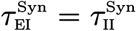 increases the level of excitation and causes the network to undergo a critical transition to an oscillatory dynamics at a Hopf bifurcation point. Consequently, larger numbers of inhibitory neurons relative to the number of excitatory neurons are required to maintain the stability of the network at lower values of 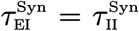. This explains the profile of the curve of Hopf points shown in Figure 7a. Disregarding the irregularly folded segment of the curve shown in Figure 7b— which can be due to the distortions observed in Figure 2b in the semi-analytically calculated response curves of excitatory neurons at low values of 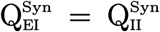—this curve of Hopf bifurcation points implies a generally similar effect when 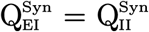 and N_i_/N are jointly varied. That is, a larger ratio of N_i_/N is required for the stability of balanced activity at lower values of 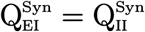. However, the sensitivity of the Hopf points with respect to changes in 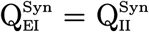 and N_i_/N appears to be particularly different. The critical (bifurcation) value of N_i_/N, below which the network transitions to oscillatory behavior, changes only slightly with sufficiently large values of 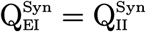. However, this critical ratio appears to be highly sensitive to excessively low values of 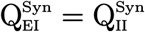.

The curve of Hopf bifurcation points shown in Figure 7c implies that both high and low values of 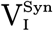 allow for network stability even at very low values of N_i_/N, although such stable activity at low values of 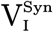 will correspond to a state of stable hyper-excitation. Moreover, it can be observed that the critical value of N_i_/N is very sensitive to both increases and decreases in 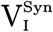 from its baseline value of −80 mV. Therefore, relatively small changes in the baseline value of 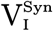 can remove the oscillatory activity of the network caused by a shortage of the number of inhibitory neurons in the network.

#### Effect of the connectivity density

The random connectivity in the local network that we study here is relatively sparse, with baseline connection probabilities P_ee_ = P_ie_ = P_ei_ = P_ii_ = 0.05 between and within excitatory and inhibitory populations. To investigate how changes in the level of sparsity in network connectivity affects the dynamic balance of excitation and inhibition, we continue the stable baseline equilibrium of the network with respect to changes in the value of overall connectivity density P_ee_ = P_ie_ = P_ei_ = P_ii_ as a codimension-one bifurcation parameter. The results are shown in Figure 8.

**Figure 8:**
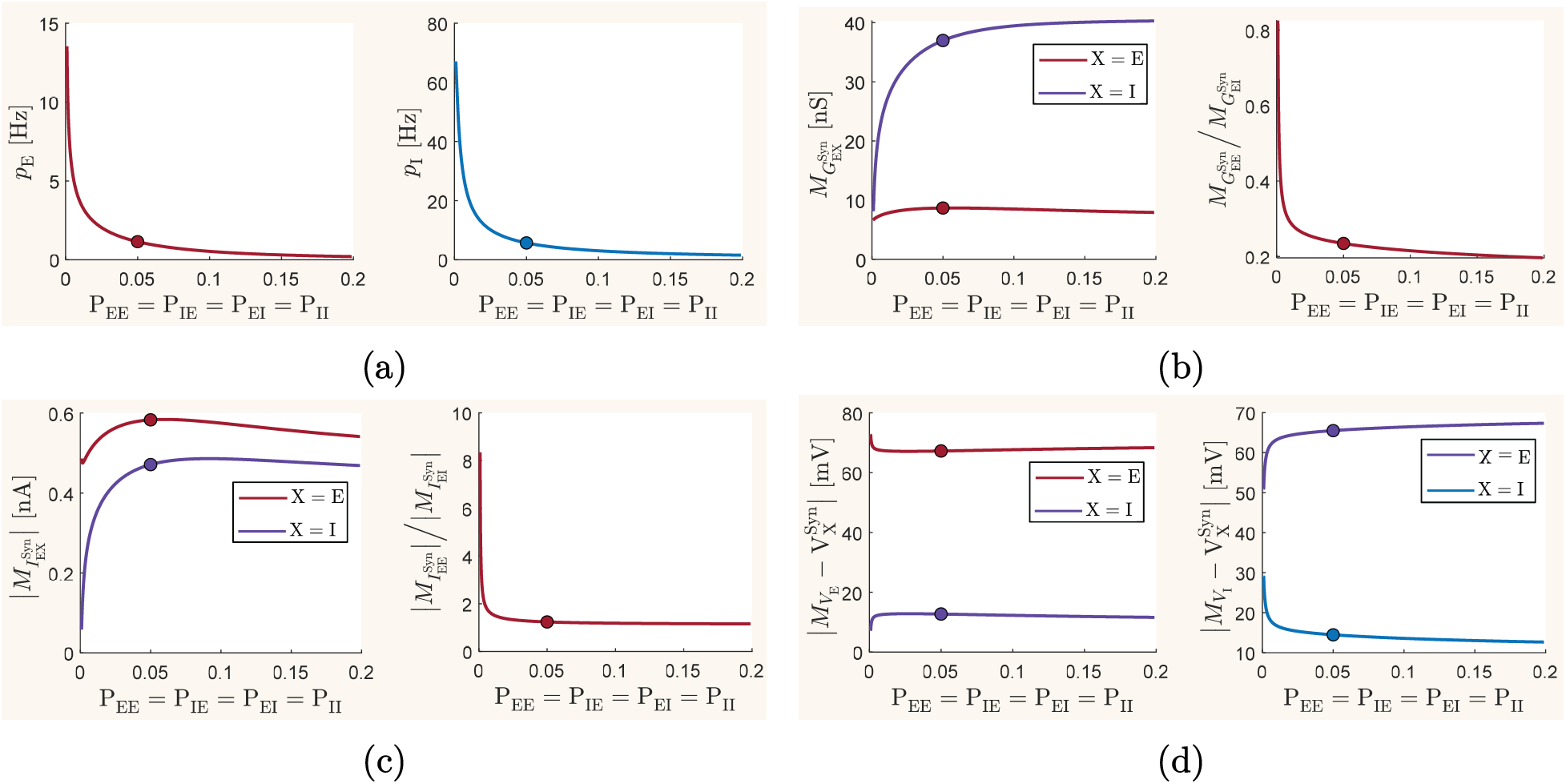
Effect of variations in the overall network connectivity density on the long-term mean-field activity of the network. All parameter values of the meanfield model, except for the connection probabilities, are set to their baseline values given in Table 1. The model is driven by background inputs of constant mean frequency 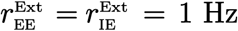. All connection probabilities are set to take the same value, which is varied as the bifurcation parameter over a biologically plausible range of values. The resulting codimension-one continuation of the network equilibrium is shown in the graphs for different network quantities. Curves of equilibria are shown by solid lines. No changes in the stability of the equilibria is detected. The points marked by dots in all graphs correspond to the baseline parameter values P_ee_ = P_ie_ = P_ei_ = P_ii_ = 0.05. We show bifurcation diagrams for (a) mean excitatory firing rate *p*_e_ and mean inhibitory firing rate *p*_i_, (b) mean synaptic conductances 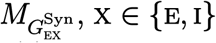, of the excitatory population, and the ratio between the two conductances, (c) absolute value of the mean excitatory and inhibitory synaptic currents, 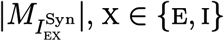, in the excitatory population, as well as the ratio between the two currents, (d) mean excitatory and inhibitory synaptic driving forces 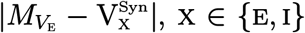, in the excitatory population, as well as such driving forces 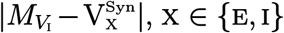, in the inhibitory population.

The mean firing rates of both excitatory and inhibitory populations decrease concurrently with increases in the overall connectivity density, as shown in Figure 8a. The ratio of the mean excitatory to inhibitory conductances shown in Figure 8b indicates that the network’s excitation-inhibition balance moves toward over-inhibition when the network’s overall connectivity density increases. On the other hand, at low values of P_ee_ = P_ie_ = P_ei_ = P_ii_, which correspond to highly sparse network connectivity, the network operates at a non-oscillatory state of hyper-excitation without undergoing a phase transition to a slow oscillatory regime. This is despite the significant decrease in mean synaptic conductances and synaptic currents at low values of overall connectivity density in the network, as it can be seen in Figures 8b and 8c. In general, we observe high sensitivity of the network balance to increases in the sparsity of the network from its baseline level, wheres much less sensitivity is observed when overall network connectivity becomes denser. Moreover—unlike the other cases that we have studied—we notice that no sustained oscillatory behavior emerges in the mean field activity of the network when over-excitation occurs due to increases in the sparsity of the network connectivity.

Besides changes in the overall network connectivity, population-specific changes in connection probabilities also affect the stability and balance of activity across the network. Recurrent excitatory-to-excitatory connections are necessary for sustaining the activity of cortical networks when they are involved in performing memory or cognitive processing tasks. Inhibitory-to-excitatory connections are crucial for stabilizing and regulating the excitatory activity in the network, and excitatory-to-inhibitory connections provide inhibitory neurons with sufficient excitation they need for their operation. However, the reason for the presence of a substantial amount of recurrent inhibitory-to-inhibitory connections in local cortical networks [54, 55], and the functionality of this type of connectivity, is less intuitive to explain. These recurrent inhibitory connections provide a level of disinhibition in the network that can regulate the overall inhibitory activity and prevent over-inhibition. However, the disinhibition created through these inhibitory-to-inhibitory connections also reduces the effectiveness of the inhibitory neurons in stabilizing the activity of excitatory neurons. To disambiguate functional implications of inhibitory-to-inhibitory connectivity, we investigate how varying only the density of this type of recurrent connectivity impacts the network balance. We choose the probability of inhibitory-to-inhibitory connections, Pii, as a bifurcation parameter and continue the baseline equilibrium state of the network. The results are shown in Figure 9.

**Figure 9:**
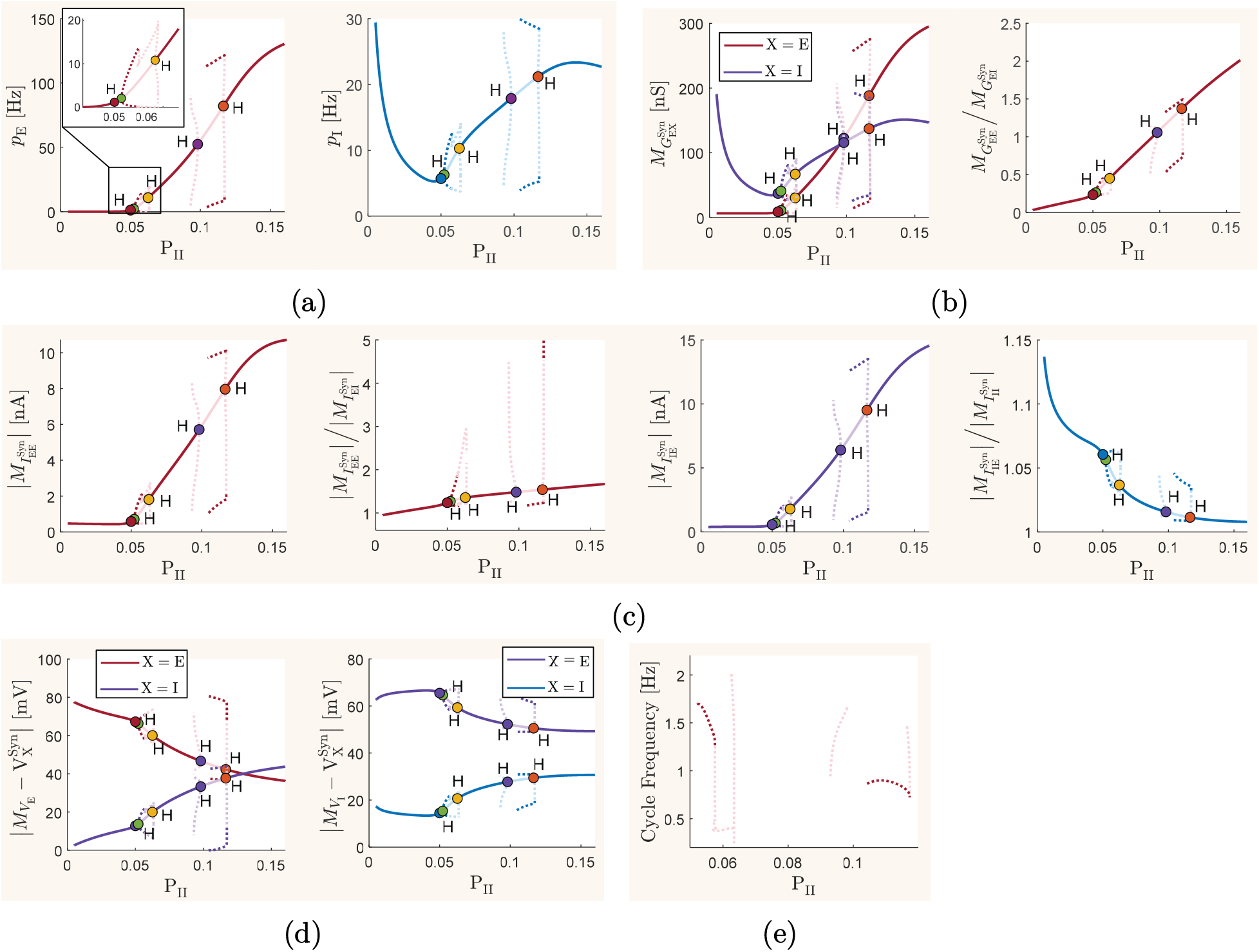
Effect of variations in the inhibitory-to-inhibitory connection probability on the long-term mean-field activity of the network. All parameter values of the mean-field model, except for the probability of connections between inhibitory neurons, are set to their baseline values given in Table 1. The model is driven by background inputs of constant mean frequency 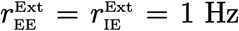. The same description as given for graphs (a)-(e) in Figure 3 holds for the graphs presented here, with the only difference being that here the bifurcation parameter for codimension-one continuation is P_ii_, with the baseline value of 0.05.

The steady-state mean firing rate curves shown in Figure 9a confirm a significant rise in the firing rate of inhibitory neurons when the level of disinhibition in the network decreases substantially at low values of P_ii_. The resulting intensive inhibition significantly hyperpolarizes the excitatory neurons, as seen in Figure 9d, and suppresses their firing activity. This shift to hyper-inhibition in the network balance reflects clearly on the steady-state ratios 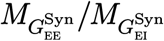 and 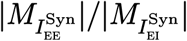, as shown in Figures 9b and 9c. In the opposite direction, however, the excessive disinhibition that results from increasing P_ii_ above its baseline value yields a relatively sharp increase in the level of excitation in the network. Excitatory neurons become exceedingly depolarized and their firing rates increases significantly. This substantial rise in the activity of excitatory neurons also depolarizes the inhibitory neurons and results in a concurrent rise in their firing rate. However, as implied from Figures 9a and 9d, the strong recurrent inhibition that is generated within the inhibitory population at large values of P_ii_ significantly restricts the mean depolarization level and firing rate of inhibitory neurons. This restrains the inhibitory neurons from effectively controlling the excitatory activity in the network, so that the excitation-inhibition balance moves to excessive excitation with increases in P_ii_. The steady-state value of the ratio 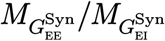 shown in Figure 9b precisely indicates such a shift in the network balance.

The bifurcation diagrams of Figure 9 further reveal that, as P_ii_ increases from its baseline value, the network’s dynamics undergoes several Hopf bifurcations, at which the stability of the network equilibrium switches and sustained oscillations emerge, or vanish. In particular, stable oscillations of delta-band frequency emerge when the density of recurrent inhibitory connectivity increases slightly above its baseline value. However, these oscillations disappear for moderately higher values of recurrent inhibitory connectivity, at which a non-oscillatory state of hyper-excitation is present across the network. The results also predict the emergence of very-high amplitude stable oscillatory activity at disproportionately large values of P_ii_. However, it should be noted that, due to the Markovian assumption used in the derivation of the mean-field model, the accuracy of the mean-field model degrades at the high firing activity associated with these oscillations. Note also that, similar to the results presented before, the curves of limit cycles originating from each Hopf bifurcation points have been continued only up to the point where the mean-field model remains mathematically valid. This numerical termination point in the continuation of limit cycles usually occurs at parameter values for which the quantity under the square root in (9) becomes negative.

### Instantaneous correlation of excitation and inhibition

The results presented so far have been obtained based on the analysis of long-term dynamics of the mean-field model, which provides information on the overall level of excitation-inhibition balance throughout the network, and how such balance is affected by changes in each of the physiological and structural parameters of the network. In experimental studies, a balance between excitation and inhibition in local cortical networks is sometimes deduced from tight correlations between the instantaneous activities of the inhibitory and excitatory populations [6,7,10]. Therefore, here we investigate how two distinct states of network activity that we observed in the results presented above, namely, a stable balanced state and a (functionally imbalanced) oscillatory state, reflect on the instantaneous correlation between the mean firing rates of excitatory and inhibitory populations.

We use the results we obtained above when we analyzed the effect of synaptic decay time constants. We solve the equations of the mean-field model corresponding to two different states: the baseline balanced state with 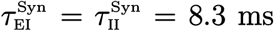, and a high-amplitude slow oscillatory state with 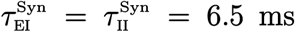. Note that the emergence of stable delta-band oscillations in the later state is ensured, as an inhibitory synaptic decay time constant of 6.5 ms is smaller than the critical Hopf bifurcation value of 7.06 ms obtained in the bifurcation analysis results shown in Figure 3. Therefore, with a constant background drive of 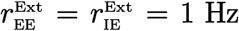, the solutions of the model with 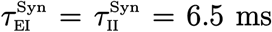 oscillate on a limit cycle, as shown in the top graph in Figure 10b.

**Figure 10:**
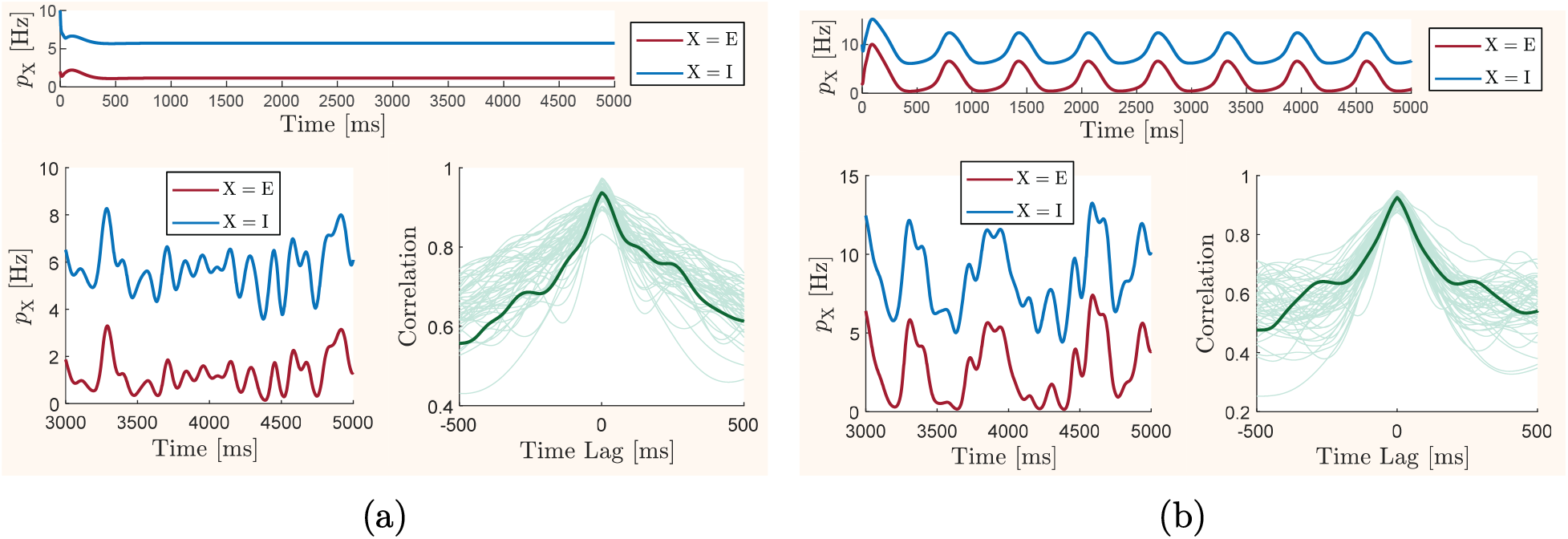
Instantaneous correlation between mean-field excitatory and inhibitory activity at baseline (balanced) and oscillatory states. At the top of each panel, instantaneous profiles of mean excitatory and inhibitory firing rates, *p*_e_ and *p*_i_, computed for the mean-field model with the constant background drive 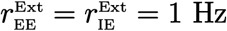, are shown over a simulation time interval of 5000 ms. All biophysical parameters of the model take their baseline values, except for 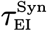 and 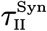 which take different values in each panel. The bottom right graph of each panel shows normalized cross-correlations between mean excitatory and inhibitory firing rates over the simulation time interval of [3000, 5000] ms. In each of these graphs, 50 correlation curves are shown that are obtained from simulating the model with 50 different background-level external drives. These external drives are generated by adding random fluctuations of amplitude 0.4 to the baseline background value of 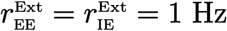. The temporal patterns of the fluctuations are generated by low-pass filtering 50 stochastic signals, each of which taking uniformly distributed random values at each instances of time. The passbands of the low-pass filters take 50 different values in the range from 1 Hz to 20 Hz. In each cross-correlation graph, a sample curve corresponding to input fluctuations filtered at passband frequency of 10 Hz is highlighted. The instantaneous profiles of the mean firing rates corresponding to the highlighted cross-correlation curves are shown on the bottom left side of each panel. (a) Instantaneous correlation between excitatory and inhibitory firing rates at the baseline balanced state, with baseline values of inhibitory synaptic decay time constants 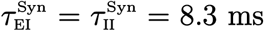. (b) Instantaneous correlation between excitatory and inhibitory firing rates at an oscillatory state emerged in the dynamics of the model when the values of inhibitory synaptic decay time constants are reduced to 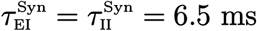.

Since a mean-field model represents network activity only at the mean-field level, it cannot reproduce the self-generated small random fluctuations typically observed in the mean firing rate or mean membrane potential of balanced *in vivo* cortical networks or balanced *in silico* networks of spiking neurons. Therefore, with a background input of constant frequency, the mean-field solutions of the model in a balanced state quickly converge to constant values, as seen in our simulation result shown in the top graph of Figure 10a. When the network transitions to an oscillatory regime, shown in the top graph of Figure 10b, the mean-field solutions with constant inputs converge to plain oscillations without any random fluctuations. The presence of tight instantaneous correlations between excitatory and inhibitory firing rates in these plain solutions is then trivial, and not very informative. Hence, in order to make informative observations, we induce random fluctuations in the mean-field solutions of the model using background inputs of randomly fluctuating frequency, instead of the constant frequency we considered for the baseline network. For each of the network states described above, we generate 50 different solution curves for *p*_e_ and *p*_i_ by driving the model with 50 different external excitatory inputs 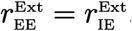. The temporal profile of these inputs is generated by adding random fluctuations of amplitude 0.4 Hz to the constant baseline inputs 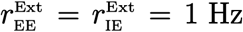. The frequency bandwidth of these fluctuations ranges approximately from 1 to 20 Hz, as described in Figure 10. To remove the effect of initial transient responses of the network, we use only the last 2 seconds of the computed solution curves for our correlation analysis described below.

A sample temporal profile of *p*_e_ and *p*_i_ is shown in Figure 10 for each state, along with normalized cross-correlations between *p*_e_ and *p*_i_ for all 50 pairs of simulated mean firing rates. In both states, the sample temporal curves show a tight instantaneous correlation between the mean excitatory and inhibitory firing rates. The normalized cross-correlation curves confirm the presence of such tight correlations in all 50 firing rate profiles. The time lags between the mean excitatory and inhibitory firing rates in both states appear to be very small: 0.50 ± 1.08 ms at the baseline balanced state, and 0.35 ± 0.74 ms at the oscillatory state, meaning that on average the inhibitory firing activity is slightly ahead of the excitatory firing activity. It should be noted though, that these tight correlations between instantaneous excitatory and inhibitory activities exist at both of the two distinct stable and oscillatory network states that we analyzed here, despite the essentially different dynamic behavior and conditions of overall excitation-inhibition balance that we observed at these states through our bifurcation analysis.

### Balanced and oscillatory activity in spiking networks

We performed our extensive bifurcation analyses presented above by taking the advantage of the simplicity and computational tractability of the mean-field model, which allowed for investigating changes in the long-term dynamics of the underlying local cortical network over a wide range of variations in the key physiological and structural parameters of the network, thereby observing the impacts of these factors on the overall balance of excitation and inhibition in the network. However, the inevitable simplifying assumptions, such as a Markovian assumption, the and semi-analytic derivations that were used in developing the mean-field model we employed in our study can raise concerns about the reliability of our predictions made based on this model. To address such concerns, to some extent, we use the network of 10,000 spiking AdEx neurons described in the Model Description section to partially verify the predictions made by the mean-field model.

Specifically, we test whether the results we obtained from the mean-field model on the state transition caused by changes in inhibitory synaptic decay time constants, as well as those obtained on instantaneous correlations between excitatory and inhibitory mean firing rates, can be replicated by a sufficiently large network of spiking neurons. We simulate the spiking neuronal network (21)-(22) at the same two dynamic states described in the results shown in Figure 10, namely, the state of balanced asynchronous irregular activity with the baseline parameter values 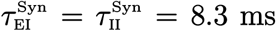, and the oscillatory state associated with reduced inhibitory decay time constants 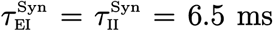. The rastergram of spiking activity at each state is shown in Figure 11 for ten percent of inhibitory and excitatory neurons.

**Figure 11:**
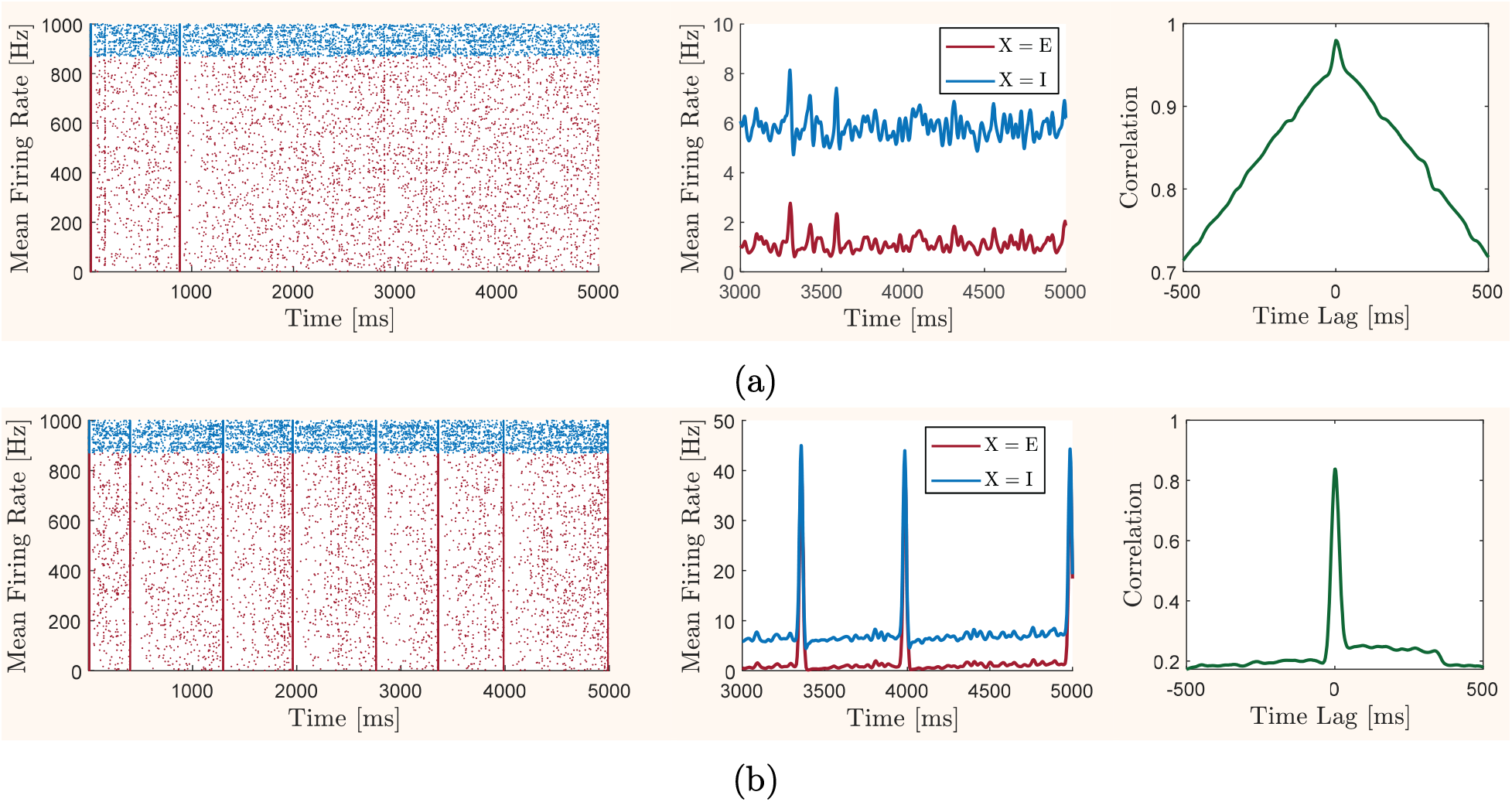
Instantaneous correlation between the average spiking activity of excitatory and inhibitory neurons at asynchronous irregular (baseline balanced) and oscillatory bursting states. In each panel, a rastergram of the excitatory (red) and inhibitory (blue) spiking activity in the spiking neuronal network (21)-(22) is shown on the left. For visual clarity, only the activity of a randomly selected 10 percent subset of total neurons are illustrated. The biophysical parameters of the spiking network are set as described in the Model Description section, except for the inhibitory synaptic decay time constants 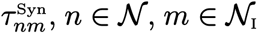 which take different values in each panel. The network receives Poisson-distributed background spike trains generated according to the description provided in the Model Description section. The middle graph in each panel shows the average firing rate of the excitatory and inhibitory neurons over the last 2 seconds of the simulation time. The graph of normalized cross-correlation between these instantaneous mean firing activities is shown on the right side of each panel. (a) Spontaneous activity and correlation between average excitatory and inhibitory firing rates at the baseline asynchronous irregular state with baseline values of inhibitory synaptic decay time constants, 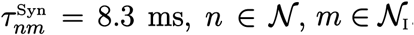. (b) Spontaneous activity and correlation between average excitatory and inhibitory firing rates at an oscillatory bursting state emerged as a result of a reduction in the values of inhibitory synaptic decay time constants to 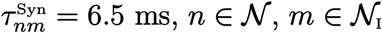.

The rastergram shown in Figure 11a demonstrates asynchronous and irregular activity at the baseline state. The mean firing rates of the excitatory and inhibitory neurons, calculated using the last 2 seconds of the spike trains in order to remove the effect of initial transient activity, are 1.13 Hz and 5.84 Hz, respectively. These rates of firing activity almost precisely match the mean firing rates *p*_e_ = 1.15 Hz and *p*_i_ = 5.71 Hz obtained at the baseline balanced equilibrium of the mean-field model. Consistent with the predictions made through bifurcation analysis of the mean-field model, the rastergram shown in Figure 11b demonstrates the emergence of a slow oscillatory bursting state in the activity of the spiking neurons when inhibitory decay time constants are reduced to 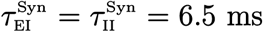, a value below the Hopf bifurcation value shown in Figure 3. Although the population bursts in the rastergram do not appear completely periodically, partly because of the stochastic nature of the Poisson-distributed input spike trains, counting the total number of bursts appearing in the rastergram of Figure 11b confirms a close agreement between the frequency of oscillatory bursting observed here with that of the mean-field oscillations shown in Figure 10b. Importantly, the spiking activity we obtained here, both at the asynchronous irregular regime and at the oscillatory bursting regime, are also closely comparable with those obtained from *in vitro* experiments and detailed *in silico* reconstruction of the rat neocortical microcircuitry, as shown in [37, Figure 12].

Similar to the results shown in Figure 10, the instantaneous profiles of mean firing rates and the normalized cross-correlation curves shown in Figure 11 indicate a tight temporal correlation between the excitatory and inhibitory spontaneous activity, in both asynchronous irregular and oscillatory bursting regimes. Consistent with the results obtained using the mean-field model, the time lag between the mean excitatory and inhibitory spiking rates obtained here are very small: 0.78 ms at the asynchronous irregular state, and 0.76 ms at the oscillatory bursting state. At both states, the spontaneous inhibitory activity appears to be slightly ahead of the excitatory activity.

## Discussion

By leveraging the computational tractability of a biologically reasonable conductance-based mean-field model, we conducted a fairly comprehensive study of how variations in some of the main synaptic and structural parameters of a local cortical network—specifically characterized by the properties of the mouse and rat neocortical microcircuitry—influence the balance between overall excitation and inhibition in the network. For this, we performed bifurcation analyses of the baseline balanced equilibrium state of the mean-field model with respect to variations in each of the key parameters of the network, namely, the synaptic (physiological) parameters such as synaptic decay time constants, synaptic quantal conductances, and synaptic reversal potentials, as well as the structural parameters such as the ratio of the number of inhibitory neurons to the total number of neurons, the density (or sparsity) of the overall network connectivity, and the inhibitory-to-inhibitory connection probability. Additionally, we used a sufficiently large network of spiking neurons to test the reliability of the predictions made based on the mean-field model. Below, we summarize and discuss the key observations of our study.

### Continuous quantification of the level of balance

The level of overall excitation and inhibition in a network is often identified in previous studies as either balanced or imbalanced. However, the degree by which the overall balance of excitation and inhibition— as we qualitatively interpreted as an operating set point for a normally functioning cortical network—deviates from its perfect level is continuously quantifiable. That means, we can quantify how balanced the network is, or how far the network activity is deviated from the balanced state. We performed such continuous quantification in our analysis. We first identified a baseline (well-) balanced reference state for the mean-field activity of our model by analyzing its long-term behavior in response to different levels of external excitatory inputs. Then, we investigated how this established balanced state is altered by changes in different parameters of the network; see Figure 1. Our results confirm that the ratio of the mean excitatory to the mean inhibitory synaptic conductances in the network—in reference to its value at the baseline balanced state—is a reliable measure for continuously quantifying the level of overall excitation-inhibition balance. Mean excitatory and inhibitory conductances can be experimentally measured, both *in vitro* and *in vivo* [56], and the ratio between them has been shown to remain constant in well-balanced states [3–5, 11, 31].

### Neuronal gain and excitability modulation

Increasing the decay time constant and the quantal conductance of inhibitory synapses decreases the gain (slope of the response curve) and excitability (horizontal shift of the response curve) of the neurons. Opposite effects are observed when the decay time constant and the quantal conductance of excitatory synapses are increased. Increases in excitatory synaptic reversal potentials result in enhancements in the gains of the neurons. Changes in the inhibitory reversal potentials have non-monotonic effects on the gain of the neurons; see Figure 2.

### Effect of synaptic parameters

Increasing the value of the three main inhibitory synaptic parameters that we studied here, to values above their baseline, moves the excitation-inhibition balance toward over-inhibition. On the other hand, decreasing these parameters below their baseline value moves the balance toward over-excitation. Modulations of the excitatory synaptic parameters have the opposite effects on the network balance, but with slightly less critical impacts on the network stability; see Figures 3–5.

### Effect of the inhibitory proportion of neurons

Changes in the ratio of the number of inhibitory neurons to the total number of neurons affect the excitation-inhibition balance in essentially similar way to how changes in inhibitory synaptic decay time constants do; see Figure 6.

### Effect of the overall network connectivity density

Increasing the sparsity of overall network connectivity moves the network balance toward over-excitation. When the network is very sparsely-connected, a further increase in the sparsity of the network connectivity causes a very sharp rise in the level of overall excitation, yet the network dynamics does not transition to an oscillatory regime. Increasing the density of overall network connectivity, on the other hand, slowly shifts the network balance to over-inhibition; see Figure 8.

### Transition to slow oscillatory regimes

In most cases in our results, sufficiently large deviations of the network balance toward over-excitation transition the network activity to an oscillatory regime. The only exception we observed was the case where we increased the overall sparsity of the network connectivity, which resulted in hyper-excited yet stable non-oscillatory states of network activity. In all cases, the emerging oscillations are slow, with their frequency being in the delta band. In particular, network oscillations emerging due to a reduction in the inhibitory proportion of the total network population or in the decay time constant of inhibitory synapses, as well as those emerging due to higher density of recurrent inhibitory-to-inhibitory connectivity, are of high amplitude; see Figures 3–5, 6, 8, and 9. High-amplitude delta oscillations are strongly correlated with loss of consciousness, and are frequently observed in states such as coma, anaesthesia, generalized epileptic seizures, and slow wave sleep [17, 57]. Therefore, it can be implied from our results that transitions to slow oscillatory regimes due to shifts in the balance of excitation and inhibition—if occurred in a sufficiently large number of local cortical networks across the neocortex—can indeed correspond to a critical state transition in the brain to a state of diminished consciousness.

### Criticality of the inhibitory synaptic conductances and the inhibitory proportion of neurons

The baseline values for inhibitory synaptic decay time constants, and for the ratio between the number of inhibitory neurons and the total number of neurons, are critical values for network stability. At such values, the network is in the well-balanced state as characterized above. However, relatively small reductions in these values result in a transition of network activity to a slow oscillatory regime; see Figures 3 and 6. This, in particular, suggests that the typical value of the inhibitory proportion of neurons in the network or, equivalently, the typical ratio between the number of inhibitory and excitatory neurons, is almost optimal. This optimality is in the sense that a larger inhibitory proportion results in redundant inhibition and a less excitable network, and a slightly smaller inhibitory proportion results in a phase transition and, possibly, loss of functionality. Therefore, the number of inhibitory neurons, relative to the number of excitatory neurons, appears to be nearly at the minimum number required to safely stabilize the overall excitatory activity under normal conditions. This optimal value, however, is in direct correspondence with the value of inhibitory synaptic decay time constants. A larger number of inhibitory neurons is required for network stability if inhibitory synapses decay faster, and a smaller number of inhibitory neurons can maintain the stability of the network if inhibitory synaptic events last longer; see Figure 7a.

### Criticality of the density of inhibitory-to-inhibitory connectivity

The density of recurrent inhibitory-to-inhibitory connectivity, relative to the density of other types of local cortical connectivity, is a crucial factor both in controlling the level of inhibitory activity in the network and in stabilizing the excitatory activity. Our results show that the baseline value for the probability of such recurrent inhibitory connectivity is critical. A moderately lower density of inhibitory-to-inhibitory connectivity results in a rather uncontrolled rise in inhibitory activity, which moves the network balance toward hyper-inhibition and yields a non-excitable network. On the contrary, a slight increase in the density of such recurrent inhibitory connectivity results in a sharp change in the balance of excitation and inhibition toward over-excitation and, subsequently, emergence of high-amplitude delta oscillations; see Figure 9.

### Robustness of network stability to changes in quantal conductances

The stability of the network in our results is fairly robust to changes in synaptic quantal conductances. That is, only a significantly large reduction in the value of inhibitory quantal conductances, or a significantly large increase in the value of excitatory quantal conductances, can destabilize the network’s equilibrium through a Hopf bifurcation; see Figure 4. This is in fact an expedient property for the functionality of the network. The values of synaptic quantal conductances are subject to dynamic changes due to plasticity of the synapses, which is essential in learning and memory. The network would otherwise be totally dysfunctional if it could be destabilized by such level of synaptic plasticity. This robustness of the network stability to changes in synaptic quantal conductances further allows for synaptic plasticity to play a key role in fine-tuning of the excitation-inhibition balance [8,10,28,34,58].

### Joint effect of synaptic and structural parameters

Balance of excitation and inhibition is established by integrated contributions of multiple physiological and anatomical factors. We mostly analyzed the effects of these factors separately. However, our limited results on the joint effect of synaptic parameters and the relative size of the inhibitory population particularly suggest that the high-amplitude delta oscillations that emerge in a local cortical network due to a reduced number of inhibitory neurons can be effectively suppressed by a reasonable amount of increase in the decay time constant or reversal potential of the inhibitory synapses, which restores the network’s stability by transitioning its dynamics through a Hopf bifurcation; see Figure 7.

### Instantaneous correlation of excitation and inhibition

Besides our extensive investigation of the overall excitation-inhibition balance through analyzing the long-term solutions of the mean-field model, we also studied the instantaneous correlation between the spontaneous excitatory and inhibitory activity in a network, using both the mean-field model and the spiking neuronal network model. Our results reveal a tight instantaneous correlation between mean excitatory and inhibitory firing rates, with an average time lag of less that one millisecond—in both the baseline balanced and the oscillatory states; see Figures 10 and 11. The presence of such tight correlation is consistent with some experimental observations, which have confirmed tightly correlated temporal variations in the activity of excitatory and inhibitory neurons in a balanced state, with a time lag of only a few millisecond between them [6,7,11,59,60]. In our simulations, on average, the mean spontaneous activity of inhibitory neurons leads that of the excitatory neurons by a fraction of millisecond, whereas experimental studies often report inhibitory activities to lag a few millisecond behind the excitatory activity, especially for evoked activities [6, 7]. The seeming discrepancy, however, is likely because the sign of the time lag between the correlated activities can depend on the physical quantities measured (e.g., firing rates, postsynaptic potentials, postsynaptic currents, etc.) as well as specific distributions and timing of external inputs to each of the inhibitory and excitatory populations. In our simulations, we provided evenly distributed background excitatory spikes at a constant average rate to both populations.

### Insufficiency of tight activity correlation for inferring normal balance

The tight instantaneous correlation between spontaneous excitatory and inhibitory activities is maintained at different levels of excitation-inhibition balance in our simulations, both in a network operating near a stable equilibrium, and in a network that has transitioned to a slowly-oscillating (-bursting) regime; see Figures 10 and 11. The presence of such a tight temporal correlation has been confirmed experimentally at different dynamic states of spontaneous, evoked, and oscillatory network activity [6,7,54,59]. However, the functionality of a cortical network, and its overall level of excitation-inhibition balance, can change substantially at these different states. For example, the presence of high-amplitude delta oscillations in a network can be associated with a state of diminished consciousness, a state functionally distinctive from the alert state. Moreover, the level of overall excitation-inhibition balance in a network, when the network is operating at different conditions, can deviate from its normal dynamic state, either toward more inhibition or toward more excitation. Importantly, such deviations do not necessarily reflect on the instantaneous correlation between excitatory and inhibitory activities. Therefore, we argue that a tight correlation between the temporal recordings of excitatory and inhibitory neuronal activities in a network is not a sufficiently strong evidence for inferring a functional balance of excitation and inhibition in the network. As we discussed above, our results suggests that the ratio between mean excitatory and inhibitory synaptic conductances is a more reliable measure of the level of overall excitation-inhibition balance in a local cortical network, as it continuously quantifies the amount of deviations from a normally balanced state.

### Reliability of the predictions

The results we presented in this paper and the key observations we summarized above were predominantly obtained using a mean-field model, which naturally suffers from inaccuracies due to simplifying assumptions. However, the model is simple and yet maintains key biological details. This allowed us the capacity to fairly comprehensively investigate the contributions of various factors in the balance of excitation and inhibition in a local cortical network. Performing such an extensive study experimentally, or using detailed biological models, would be rather impractical. To test the reliability of the predictions made based on the mean-field model, we performed a sample study using a more detailed framework of a spiking network. The close agreement between the results we obtained from this spiking network and those predicted based on the mean-field model—which were also consistent with the spiking activity observed in *in vitro* experiments and in *in silico* computations of a detailed neocortical microcircuitry [37, Figure 12]—is promising. Hence, we anticipate that the observations we made in this paper can inform future experimental and theoretical studies on understanding the homeostatic mechanisms involved in regulation of the excitation-inhibition balance, and on identifying the pathological causes of short-term and long-term disturbances in this essential operating condition of local cortical networks.

## Funding acknowledgment

This work was supported by the National Eye Institute of the National Institutes of Health under Award Number R00 EY030840. The content is solely the responsibility of the authors and does not necessarily represent the official views of the National Institutes of Health.

## Author contribution

We use the CRediT Taxonomy to describe each author’s contributions to the work, as follows: Conceptualization (FS), Data Curation (FS), Formal Analysis (FS), Funding Acquisition (HC), Investigation (FS), Methodology (FS), Project Administration (FS, HC), Resources (N/A), Software (FS), Supervision (N/A), Validation (FS), Visualization (FS), Writing– Original Draft Preparation (FS), Writing–Review & Editing (FS, HC)

